# *C9orf72* poly(PR) mediated neurodegeneration is associated with nucleolar stress

**DOI:** 10.1101/2023.02.16.528809

**Authors:** ME Cicardi, JH Hallgren, D Mawrie, K Krishnamurthy, SS Markandaiah, AT Nelson, V Kankate, EN Anderson, P Pasinelli, UB Pandey, CM Eischen, D Trotti

**Affiliations:** Jefferson Weinberg ALS Center; Vickie and Jack Farber Institute for Neuroscience, Department of Neuroscience, Thomas Jefferson University, Philadelphia, PA, USA; Sidney Kimmel Cancer Center, Department of Pharmacology, Physiology, and Cancer Biology, Thomas Jefferson University, Philadelphia, PA, USA; Center for Neuroscience, Department of Human Genetics, University of Pittsburgh Graduate School of Public Health, Pittsburgh, PA, USA

## Abstract

The ALS/FTD-linked intronic hexanucleotide repeat expansion in the *C9orf72* gene is translated into dipeptide repeat proteins, among which poly-proline-arginine (PR) displays the most aggressive neurotoxicity *in-vitro* and *in-vivo*. PR partitions to the nucleus when expressed in neurons and other cell types. Using *drosophila* and primary rat cortical neurons as model systems, we show that by lessening the nuclear accumulation of PR, we can drastically reduce its neurotoxicity. PR accumulates in the nucleolus, a site of ribosome biogenesis that regulates the cell stress response. We examined the effect of nucleolar PR accumulation and its impact on nucleolar function and determined that PR caused nucleolar stress and increased levels of the transcription factor p53. Downregulating p53 levels, either genetically or by increasing its degradation, also prevented PR-mediated neurotoxic phenotypes both in *in-vitro* and *in-vivo* models. We also investigated whether PR could cause the senescence phenotype in neurons but observed none. Instead, we found induction of apoptosis *via* caspase-3 activation. In summary, we uncovered the central role of nucleolar dysfunction upon PR expression in the context of C9-ALS/FTD.

## INTRODUCTION

The most common genetic cause of amyotrophic lateral sclerosis (ALS) and frontotemporal dementia (FTD) is an intronic G4C2 nucleotide repeat expansion (NRE) of the C9orf72 gene (C9) (Biasiotto & Zanella, 2018; Dejesus-hernandez *et al*, 2011; Renton *et al*, 2011)’. This expansion in C9 transcripts leads to a decreased C9ORF72 protein. NRE-containing RNA transcripts can bind to multiple nuclear proteins, resulting in nuclear RNA foci (Lee *et al*, 2013). Finally, NRE-containing RNA transcripts can be exported to the cytoplasm and translated through a mechanism known as repeat-associated non-AUG translation (RAN-T), which produces 5 different dipeptide repeat proteins (DPRs) from the sense and antisense direction: poly glycine-alanine (GA), poly glycine-proline (GP), and poly glycine-arginine (GR); poly proline-alanine (PA) and poly proline-arginine (PR) (Ash *et al*, 2013). PR is the most neurotoxic among the DPRs, as shown in many cell and animal models (Jovičič *et al*, 2015; Zhang *et al*, 2019; Zhang *et al*, 2018; Zhang *et al*, 2014). A large fraction of PR accumulates in the nucleolus as it’s been demonstrated by immunofluorescence experiments and BioID labeling (Liu *et al*, 2022). Even though many pathways have been associated with PR toxicity so far, little is known about whether PR has detrimental effects on normal nucleolar functions and which nucleolar downstream pathways are affected. The nucleolus is a membrane-less subnuclear compartment formed by multiple proteins coalescing around ribosomal DNA (rDNA) sites to transcribe ribosomal RNA (rRNA) and assemble ribosomes (Boulon *et al*, 2010). It is a central hub for biological processes and a sensor of cellular stress (Lee *et al*, 2016).

In particular, the nucleolus is composed of three regions: a fibrillar core (FC), where rRNA is transcribed; a dense fibrillar component (DFC), where pre-RNA is processed; and a granular component (GC), where pre-ribosome subunits are assembled (Boisvert *et al*, 2007). Disaggregation of the nucleolus due to different types of insult precedes impaired ribosomal biogenesis and leads to cell death by apoptosis (Boisvert *et al*., 2007). P53 is the most well-known mediator of apoptosis, and its regulatory pathways have been central to numerous studies. One of the most important regulators of p53 levels is Mdm2 (Marine & Lozano, 2010). Under unstressed conditions, p53 is bound to Mdm2, which ubiquitinates p53 to target it to the proteasome. When a cellular insult occurs, such as DNA damage, translation stalling, or mitochondrial stress, a series of effectors exit the nucleolus (NPM, ARF, RPL11, RPL5) and bind Mdm2, inhibiting the ubiquitination of p53 and leading to increased p53 levels (Marine & Lozano, 2010). In alternative to triggering apoptosis, p53 can also mediate the induction of a secondary pathway known as senescence, which prolongs cell life, preserving some of the cell’s functions while delaying cell death (Jurk *et al*, 2012). Senescence is mainly mediated by the cyclin-dependent kinase inhibitors p21(CDKN1A) and p16 (CDKN2A), with recent evidence suggesting a role for transcription factors, including NF-kB (Chen & Stark, 2019; Coppé *et al*, 2014; Salminen *et al*, 2012; Van Deursen, 2014). Senescence in proliferative cells causes a halt in cell replication. Still, even though neurons are post-mitotic cells, a process termed “senescence-like” has been reported to occur in neurons, both *in-vitro* and *in-vivo* (Martínez-Cué & Rueda, 2020; Vazquez-Villasenor *et al*, 2020). Given this premise, it is straightforward to predict that changes or perturbations in the nucleolus might trigger an array of downstream events. Indeed, we hypothesized that the neuronal stress posed by PR localization to the nucleolus could change its normal functions, leading to a p53 increase and inducing either a senescence-like response in neurons or apoptosis. We showed here that PR caused nucleolar dysfunctions and increased p53 levels. We then characterized both senescence and apoptosis activation triggered by p53 accumulation in neurons. Lastly, we intervened by inhibiting the p53 activation pathway to rescue PR neurotoxicity.

## RESULTS

### PR yields nucleolar stress without influencing nucleolar dynamics

The nucleolus is a membrane-less structure characterized by three layers: fibrillar core (FC), dense fibrillar component (DFC), and granular component (GC). We stained primary rat cortical neurons transfected with PR (50 dipeptide repeats) for nucleophosmin (NPM) and nucleolar transcription factor 1 (UBF-1), proteins expressed in the GC and the FC, respectively. PR_50_ (GFP reporter tagged at the C-terminus) co-localizes with NPM and UBF1 (**Fig. 1A, EV1 A-B**), suggesting that PR_50_ is in both nucleolar layers. PR_50_-GFP is also present in the nucleoplasm, as indicated by the diffuse GFP fluorescence observed around the nucleolus (**Fig. 1A**).

**Figure 1.**
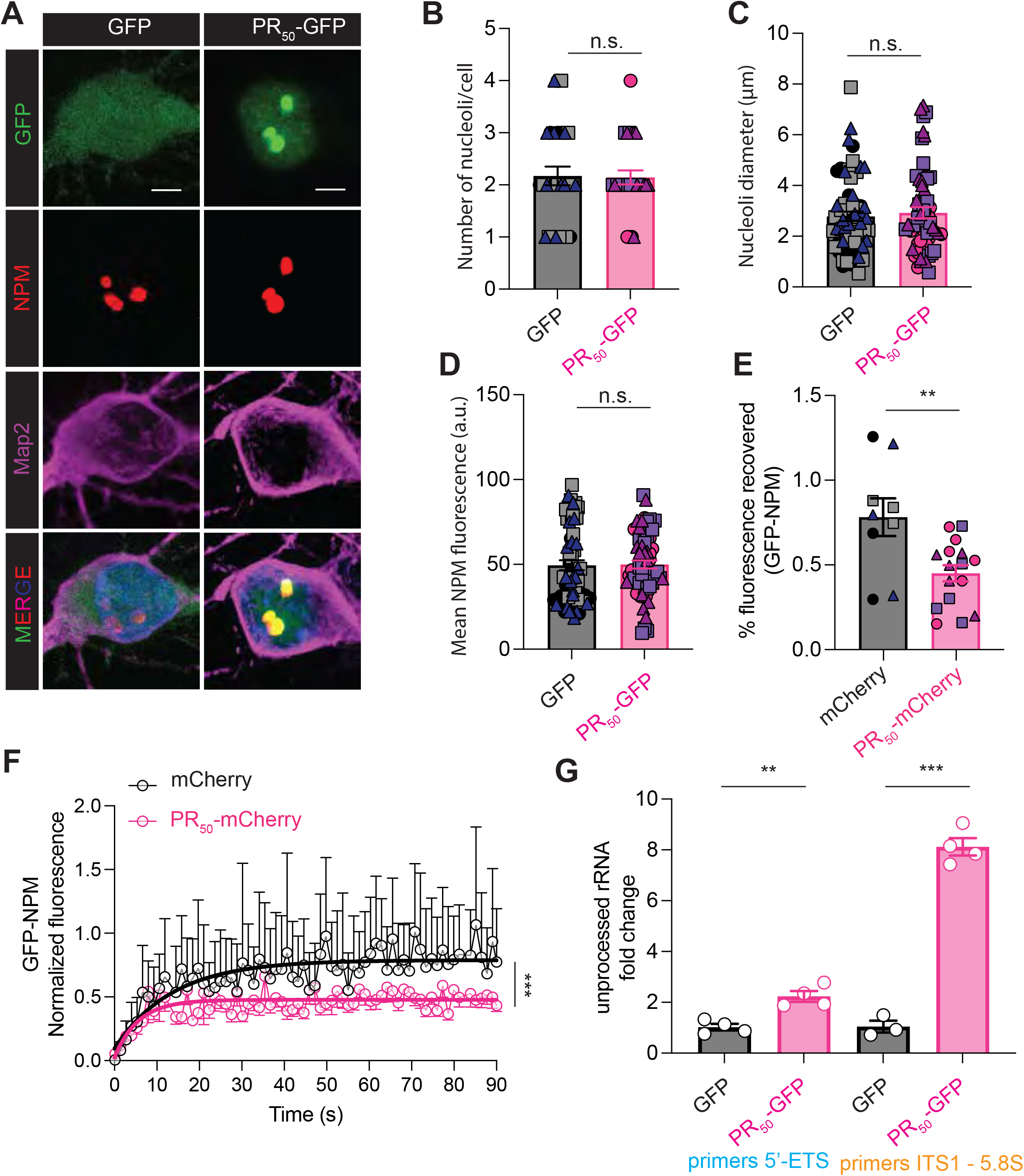
PR localization to the nucleolus induces nucleolar stress. A. Representative images of rat primary cortical neurons overexpressing GFP or PR_50_-GFP. GFP is shown in green, NPM is shown in red, Map2 is shown in magenta, and nuclei (Hoechst) are shown in blue. Scale bar = 2.5 μm B. Graph bar showing quantification of nucleoli number/cell (mean ± s.e.m., number of experiments = 3, number of neurons counted per experiment ≥10, Student’s t-test, nonparametric, p=0.8958) C. Graph bar showing quantification of nucleoli dimension (mean ± s.e.m., number of experiments = 3, number of neurons counted per experiment ≥10, Student’s t-test, nonparametric, p=0.6205) D. Graph bar showing quantification of nucleoli brightness (NPM fluorescence) (mean ± s.e.m., number of experiments = 3, number of neurons counted per experiment ≥10, Student’s t-test, non-parametric, p=9049). E. Graph bar showing quantification of FRAP percentage of recovered fluorescence of GFP-NPM after photobleaching in presence of mCherry or PR_50_-mCherry (mean ± s.e.m., number of experiments = 3, number of neurons counted per experiment ≥3, Student’s t-test, non-parametric, p=0.0042) F. Graph showing percentage of recovered fluorescence of GFP-NPM over time in presence of mCherry or PR_50_-mCherry (mean ± s.e.m., number of experiments = 3, number of neurons counted per experiment ≥3, two-way ANOVA, p<0.0001). G. qPCR of rat primary cortical neurons transfected with GFP or PR_50_-GFP for pre-processed rRNA (mean ± s.e.m., number of experiment = 4, Student’s t-test, non-parametric, ** p=0.0028, **** p<0.0001).

We then analyzed whether PR could modify the nucleolus’s morphology. In rat primary cortical neurons expressing PR_50_-GFP or GFP, we measured the number and diameter of the nucleoli and found no significant changes (**Fig. 1B-C**). NPM fluorescence intensity signal was also not different between these two groups (**Fig. 1D**).

As a membrane-less organelle, the nucleolus has liquid droplet-like properties, which can be investigated by fluorescence recovery after photobleaching (FRAP). We measured the FRAP half-time of NPM in PR_50_-GFP vs. GFP expressing neurons and found that PR_50_-GFP caused a significant decrease in the percentage of recovered fluorescence (**Fig. 1E-F, EV1 D**).

These results showed that nucleolar accumulation of PR in primary neurons did not significantly change the nucleolus morphology but affected its liquid droplet-like dynamics, in agreement with a previous study (Lee *et al*., 2016), which reported substantial changes in the liquid droplet-like properties of the nucleolus in the immortalized cancer cell lines, HEK293 and HeLa. We next checked for evidence of nucleolar stress potentially caused by PR. A well-established sign of nucleolar stress is the impaired processing of pre-rRNA to mature rRNA (Boulon *et al*., 2010). We performed qPCR on neurons expressing PR_50_-GFP or GFP as a control to quantify their respective levels of pre-rRNA (**EV1 C**). We found an increased expression of pre-processed rRNA in the presence of PR_50_-GFP, indicative of nucleolar stress (**Fig. 1G**).

### Increased expression levels of the transcription factor p53 are central to PR-mediated neurotoxicity

Activation of the p53 pathway, which is finely regulated in the nucleolus (Boulon *et al*., 2010), is the first outcome of nucleolar stress. Using confocal microscopy and immunofluorescence, we quantified p53 nuclear intensity in cells expressing either GFP or PR_50_-GFP (**Fig. 2A**). Single-cell analysis revealed increased expression levels of nuclear p53 in PR_50_-GFP expressing neurons compared to controls (**Fig. 2A, B**). We then performed western blot (WB) analysis of neurons expressing GFP and PR_50_-GFP. We also analyzed GP_50_-GFP expressing neurons as a control DPR known not to localize to the nucleolus (Wen *et al*, 2014). We observed a significant ~50% increase in p53 expression levels in PR_50_-GFP neurons compared to GFP and GP_50_-GFP (**Fig. 2C-D**). Elevated levels of p53 lead to neuronal cell death (Morrison & Kinoshita, 2000). We thus stained for cleaved caspase 3 (CC3), one of the intermediates of the p53-induced apoptotic cascade, and we found that PR_50_-GFP expressing neurons had higher levels of CC3 than GFP controls (**Fig. 2 E-F**). Through staining with lamin B, we also assessed the shape of the nucleus in the presence of PR_50_-GFP and found a decrease in the circularity score and a reduction in the fluorescence of lamin B in the presence of PR_50_-GFP (**Fig. 2 G-I**). Misshapen nuclei and loss of lamin B in neurons have been associated with neurodegeneration, further strengthening the evidence of neuronal death upon PR_50_-GFP expression (Matias *et al*, 2022).

**Figure 2.**
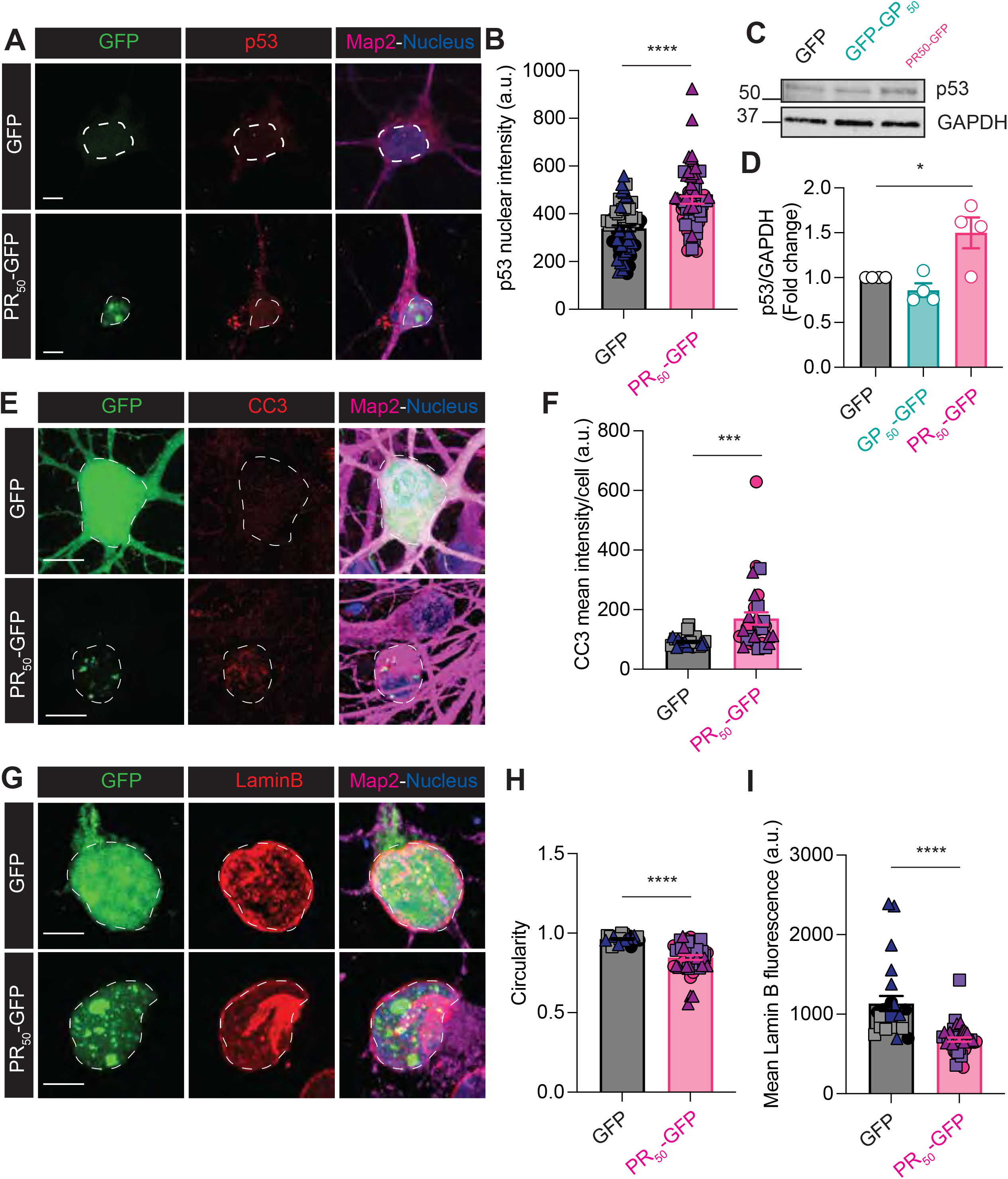
PR-mediated nucleolar stress causes activation of caspase-3 and neuronal apoptosis. A. Representative images showed rat primary cortical neurons transfected with GFP or PR_50_-GFP. GFP or PR_50_-GFP are shown in green, p53 is in red, Map2 is in magenta, and nuclei (Hoechst) are in blue. Scale bar = 10 μm B. Graph bar showing quantification of p53 nuclear intensity in rat primary cortical neurons transfected with GFP or PR_50_-GFP (mean ± s.e.m., number of experiments = 3, number of neurons counted per experiment ≥10, Student’s t-test, non-parametric, **** p < 0.0001). C. Western blot for p53 and GAPDH performed on rat primary cortical neurons transfected with GFP or GP_50_-GFP, PR_50_-GFP. D. Graph bar showing p53 quantification of Western blot. GFP vs. PR_50_-GFP * p< 0.05 (mean ± s.e.m., number of experiments = 4, Student’s t-test, non-parametric, * p = 0.0269). E. Representative images showed rat primary cortical neurons transfected with GFP or PR_50_-GFP. GFP or PR_50_-GFP are shown in green, CC3 is in red, Map2 is in magenta, and nuclei (Hoechst) are in blue. Scale bar = 10 μm F. Graph bar showing quantification of CC3 nuclear intensity in rat primary cortical neurons transfected with GFP or PR_50_-GFP (mean ± s.e.m., number of experiments = 3, number of neurons counted per experiment ≥10, Student’s t-test, non-parametric, *** p = 0.0007). G. Representative images showed rat primary cortical neurons transfected with GFP or PR_50_-GFP. GFP or PR_50_-GFP are shown in green, and Lamin B is in red, Map2 is in magenta, and nuclei (Hoechst) are in blue. Scale bar = 10 μm H. Graph bar showing quantification of nucleus circularity in rat primary cortical neurons transfected with GFP or PR_50_-GFP (mean ± s.e.m., number of experiments = 3, number of neurons counted per experiment ≥10, Student’s t-test, non-parametric, **** p< 0.0001). I. Graph bar showing quantification of LaminB nuclear intensity in rat primary cortical neurons transfected with GFP or PR_50_-GFP (mean ± s.e.m., number of experiments = 3, number of neurons counted per experiment ≥10, Student’s t-test, non-parametric, **** p< 0.0001).

### Nucleolar PR accumulation does not induce senescence

A senescence phenotype is an additional outcome of nucleolar stress and increased p53 levels in cells. To assess whether neurons expressing PR underwent senescence-like processes, we investigated whether senescence markers (Shaker *et al*, 2021) were present.

The two cyclin-dependent kinase inhibitors, p16 and p21, and the nuclear transcription factor NF-kB are the three proteins associated with senescence regulation and activation (Van Deursen, 2014). To achieve a broader expression of GFP and PR_50_-GFP, needed for WB and qPCR experiments, we transduced primary rat cortical neurons with lentivirus expressing either GFP or PR_50_-GFP and quantified the levels of these key regulators of senescence by WB. The p16, p21, and NF-kB levels were unchanged in cells expressing PR_50_-GFP compared to GFP (**Fig. 3 A-D**) at three days post-transduction. This result was also confirmed by qPCR analysis of the respective transcripts (**Fig. 3E**). Furthermore, we determined whether the cytokine transcripts that are usually increased in cells undergoing senescence (Interleukin 6 (Il-6), Interleukin (Il-1a), Interleukin (Il-1b)) were upregulated. Still, we found no induction of these transcripts (**Fig. 3E**). We also measured Il-6 released in the media by ELISA and did not detect a change between PR_50_-GFP and GFP transduced neurons (**Fig. 3F**). Similarly, the activity of lysosomal β-galactosidase, a hydrolase that converts galactose into glucose and a common feature of senescent cells, is unchanged (**Fig. 3G-H**).

**Figure 3.**
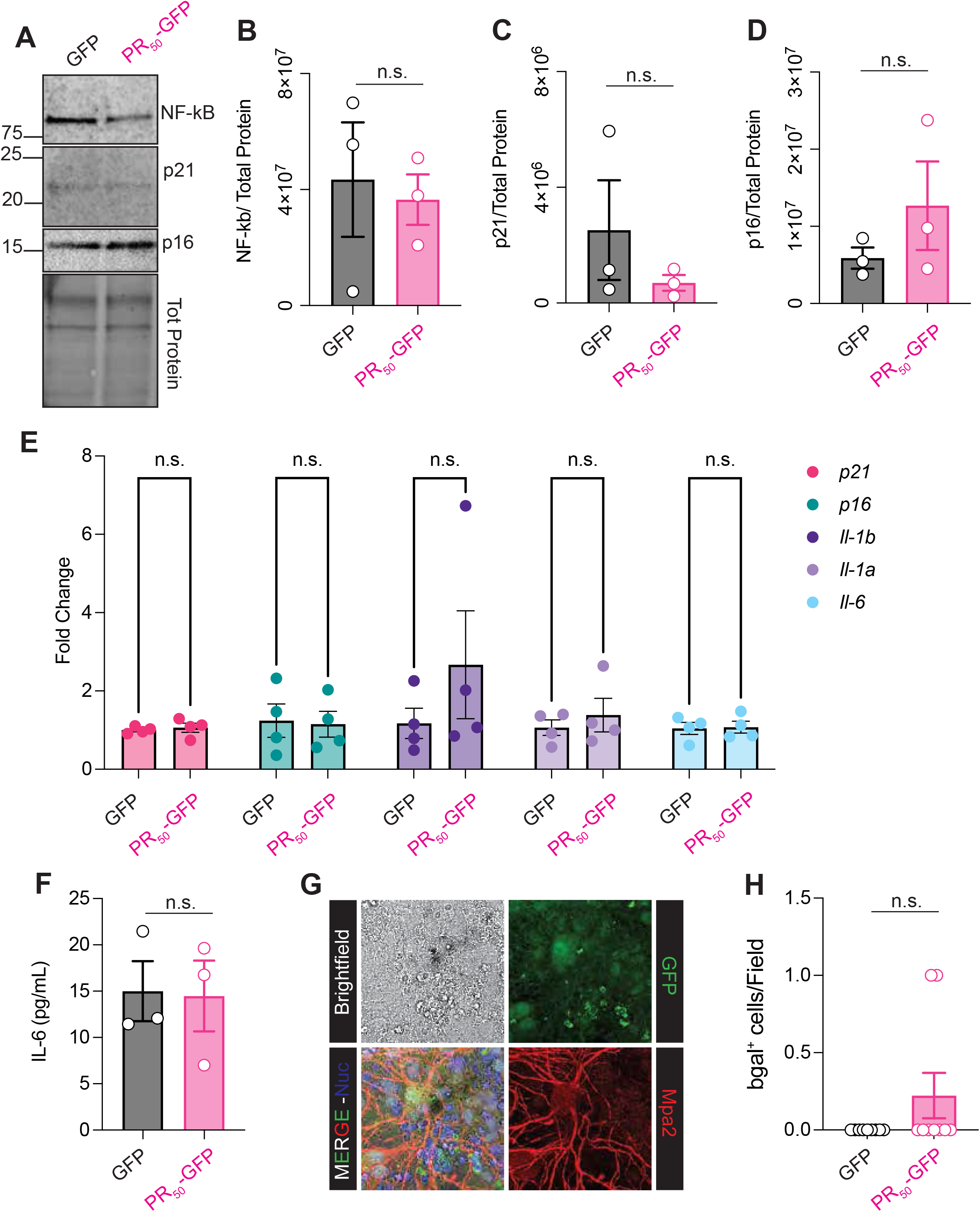
Absence of the senescence phenotype in PR-expressing neurons. A. Western blot for p21, p16, and NF-kB and total protein performed on primary cortical neurons transduced with GFP or PR_50_-GFP, 3 days post-transduction. B. Graph bar showing NF-kB quantification of Western blot performed on primary cortical neurons transduced with GFP or PR_50_-GFP (mean ± s.e.m., number of experiments = 3, Student’s t-test, non-parametric, p = 0.7654). C. Graph bar showing p21 quantification of Western blot performed on primary cortical neurons transduced with GFP or PR_50_-GFP (mean ± s.e.m., number of experiments = 3, Student’s t-test, non-parametric, p = 0.3553). D. Graph bar showing p16 quantification of Western blot performed on primary cortical neurons transduced with GFP or PR_50_-GFP (mean ± s.e.m., number of experiments = 3, Student’s t-test, non-parametric, p = 0.3144). E. Graph bar showing quantification of qPCR for *p21, p16, Il6, Il1a*, and *Il1b* performed on primary cortical neurons transduced with GFP or PR_50_-GFP, 3 days post-transduction (mean ± s.e.m., number of experiments = 3, two-way ANOVA, Sidak’s multiple comparison test, p = n.s.). F. ELISA assay for Il-6 concentration in cell culture media from primary cortical neurons transduced with GFP or PR_50_-GFP, 3 days post-transduction (mean ± s.e.m., number of experiments = 3, Student’s t-test, non-parametric, p = 0.9220). G. Representative image of positive controls for β-galactosidase staining in neurons. Positive cells are circled in green; negative cells are circled in black. Scale bar = 10μm H. Senescence-associated β-galactosidase staining was performed on primary cortical neurons transduced with GFP or PR_50_-GFP, 3 days post-transduction (mean ± s.e.m., number of experiments = 3, Student’s t-test, non-parametric, p = 0.1501).

Finally, we reasoned that the onset of the senescence phenotype could occur at a delayed time post-PR_50_ expression. To determine whether induction of senescence was instead happening in the surviving PR-expressing neuron population at later times, we measured the levels of senescence-associated markers (p16, p21, NF-kB) 7 days after PR transduction, and no significant increase by WB in any of them. qPCR again did not detect an increase in any of the senescence-associated marker transcripts (**EV2 A-E**).

Additionally, ELISA for IL-6 did not show differences between GFP and PR_50_-GFP (**EV2 F**). These data indicate that PR-expressing neurons do not undergo senescence but rather undertake the apoptosis pathway to cell death.

### Nuclear localization is required for PR-mediated neurotoxicity

PR is known to partition in the nucleus of cells, explicitly localizing to the nucleolus, and to cause fast and aggressive neuronal death(Wen *et al*., 2014). To investigate whether PR nuclear partitioning is a requirement for neurotoxicity, we heterologously expressed PR in cultured rat cortical neurons using a construct that has 3 nuclear export sequences fused to the N-terminus of the PR encoding sequence (PR_50_-GFP-3xNES) and one encoding just the PR_50_ dipeptide with no nuclear partition tag linked to it. We showed by immunofluorescence that the triple nuclear export tag added to the PR_50_ sequence significantly diminished its co-localization to the nucleolar-specific marker protein NPM (**Fig. 4A-B**). To rule out a possible difference in expression between PR_50_-GFP and PR_50_-GFP-3xNES, we measure RNA levels by qPCR and protein levels by WB of the two constructs: we found no significant differences in the expression levels of PR_50_-GFP and PR_50_-GFP-3xNES (**Fig 4C-E**), confirming that the decreased localization to the nucleolus is dependent on the presence of the 3XNES and not to a different expression rate.

**Figure 4.**
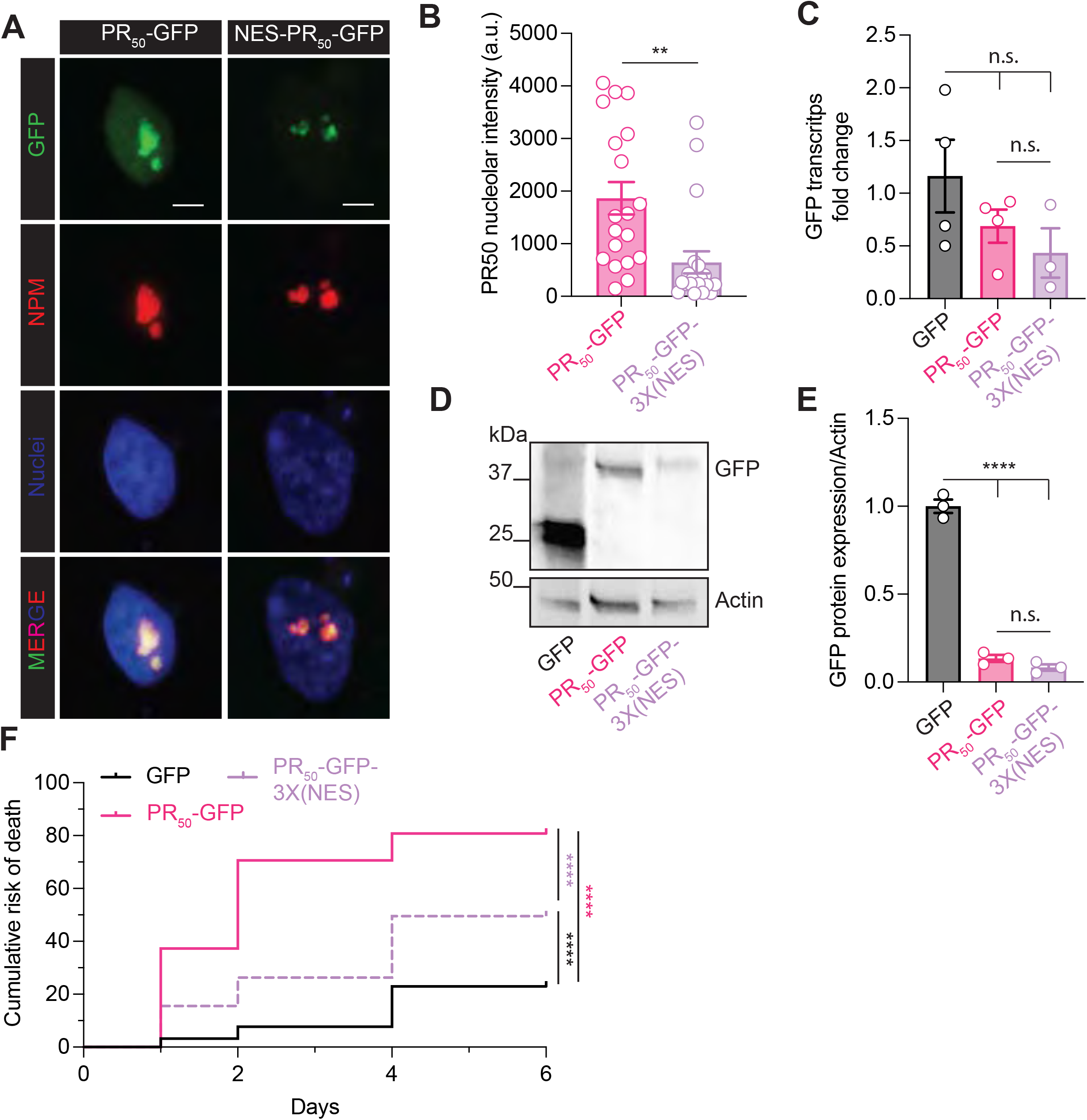
PR neurotoxicity depends on its nucleolar localization. A. Representative images of rat primary cortical neurons overexpressing PR_50_-GFP and PR_50_-3xNES-GFP. GFP is shown in green, NPM is shown in red, Map2 is shown in magenta, and nuclei (Hoechst) are shown in blue. Scale bar = 2.5 μm B. Graph bar showing quantification of GFP intensity inside nucleoli (mean ± s.e.m., number of experiments = 3, number of neurons counted per experiment ≥6, Student’s t-test, nonparametric, ** p = 0.0021). C. Graph bar showing quantification of qPCR for *GFP* performed on rat primary cortical neurons transfected with GFP or PR_50_-GFP or PR_50_-3xNES-GFP (mean ± s.e.m., number of experiments = 3, One-way ANOVA, Tukey’s multiple comparison test, p = n.s.). D. Western blot for GFP and Actin performed on rat primary cortical neurons transfected with GFP or PR_50_-GFP or PR_50_-3xNES-GFP. E. Graph bar showing GFP quantification of Western blot performed on rat primary cortical neurons transfected with GFP or PR_50_-GFP or PR_50_-3xNES-GFP. P=n.s., **** p<0,0001 (mean ± s.e.m., number of experiments = 3, One-way ANOVA, Tukey’s multiple comparison test, **** p < 0.0001). F. Kaplan Meier survival analysis curves of primary rat cortical neurons transfected with the indicated constructs (number of experiments = 3, number of neurons counted per experiment ≥100, Mantel-Cox test, details in table 1–2).

Using GFP-expressing neurons as a reference group (**Fig. 4F**), we followed the transfected cortical neurons by live-cell imaging microscopy to assess their intrinsic risk of death over several days *in vitro*. We found that PR_50_ decreased neuron viability compared to GFP within 24 hours (p<0.0001, hazard ratio of PR_50_-GFP vs. GFP = 10.32, **Table 1, Fig. 4F**). The fusion of a 3xNES tag to the PR_50_ expressing construct (PR_50_-GFP-3xNES) significantly increased neuronal viability (p<0.0001, hazard ratio of PR_50_-GFP vs. PR_50_-GFP-3xNES neurons = 3.187, **Table 1**, **Fig. 4F**), suggesting that the levels of PR localizing in the nucleolus are critical to inducing apoptosis.

**Table 1:** Mantel-Cox test summary for neuronal survival data of GFP vs PR50 relative to Figure 4

| GFP vs PR50 |  |  |
| --- | --- | --- |
| Log-rank (Mantel-Cox) test |  |  |
| Chi square | 287 |  |
| df | 1 |  |
| P value | <0.0001 |  |
| P value summary | **** |  |
| Are the survival curves sig different? | Yes |  |
| Hazard Ratio (Mantel-Haenszel) | GFP vs PR50 | PR50 vs GFP |
| Ratio (and its reciprocal) | 0.09687 | 10.32 |
| 95% CI of ratio | 0.07394 to 0.1269 | 7.880 to 13.52 |

We then sought confirmation of these results in an organismal model. PR_50_-GFP caused robust retinal degeneration compared to GFP or GFP-NES when expressed in the *Drosophila* eye (**EV3 A-B**). Notably, flies overexpressing PR_50_-GFP-NES showed a 50% reduction of the eye degeneration score (**EV3 B**), supporting the notion that PR exerts toxicity when localized to the nucleus. By immunofluorescence we then observed that in the flies eyes the nuclear localization of PR_50_-GFP is significantly reduced when the NES signal is present (**EV3 C-D**).

### Modulating p53 levels rescues PR toxicity

We hypothesized that modulating P53 levels could be a valuable strategy to abate PR neurotoxicity. To test this, we first co-expressed a dominant-negative form of p53 in primary cortical neurons to rescue *in vitro* viability of neurons expressing PR_50_-GFP. We followed neurons transfected with PR_50_-GFP or GFP for seven days by live imaging. As we previously showed, over seven days, we observed that PR_50_-GFP caused pronounced toxicity in cortical neurons that were significantly rescued by co-expression of the dominant-negative form of p53 (p53-DN) (**Fig. 5A, Table 2**).

**Figure 5.**
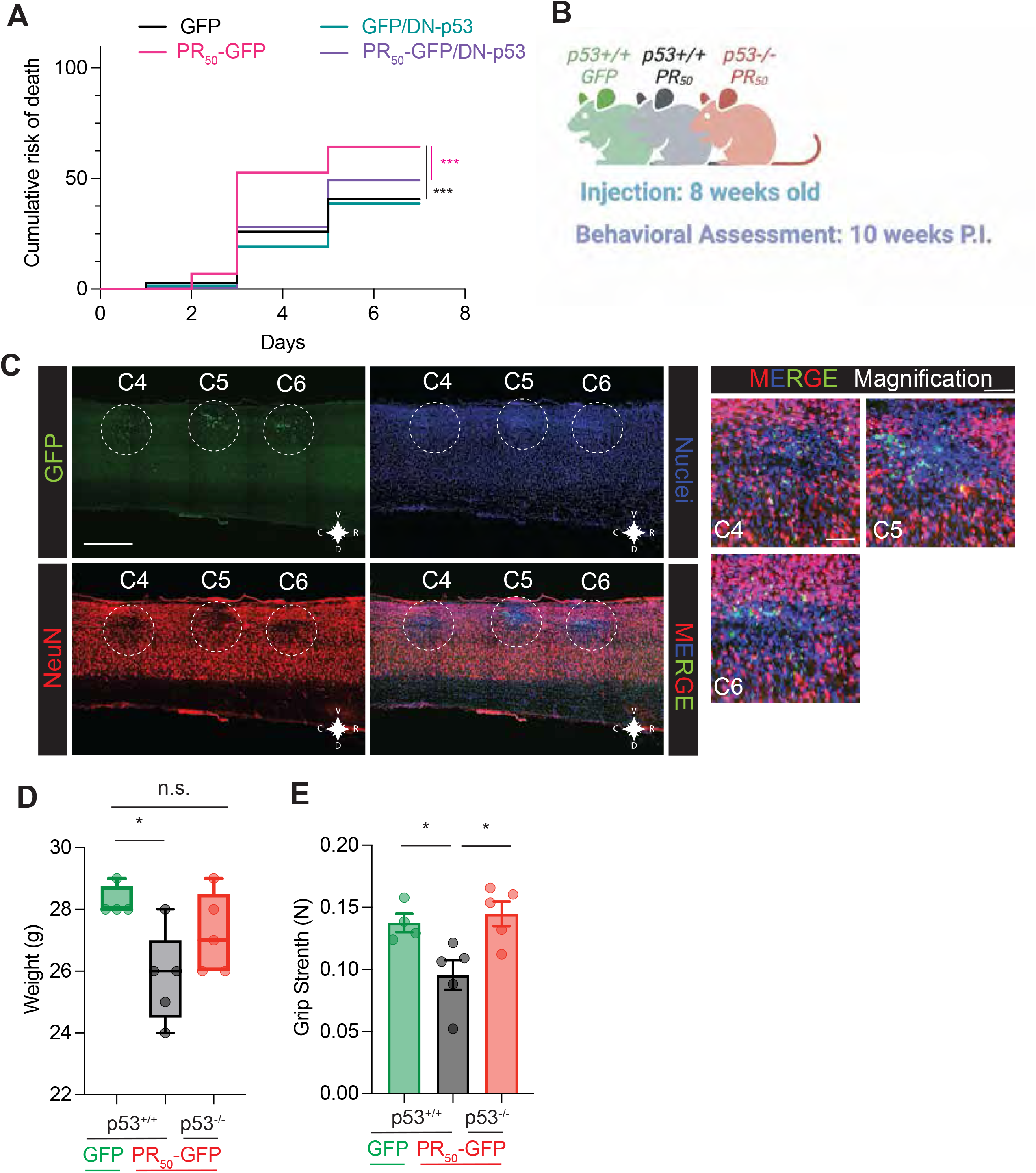
PR-mediated neurotoxicity is partially rescued by manipulating the p53 pathway. A. Kaplan-Meier survival analysis curves of primary rat cortical neurons transfected with GFP or PR_50_-GFP and pcDNA3 or DN-p53 (dominant negative). *** p< 0.001 (number of experiments = 3, number of neurons counted per experiment ≥100, Mantel-Cox test, details in table 3–4). B. Schematic and timeline of the experiment. C. Left panel: Representative images of a sagittal section of the spinal cord showing the three injection sites: GFP is in green, NeuN is in red, and nuclei are in blue (Hoechst). Scale bar= 500μm. C= caudal, V= ventral, R= rostral, D= dorsal. Right panel: Enlargement of each injection site. GFP is shown in green, NeuN is shown in red, and nuclei are shown in blue (Hoechst). Scale bar= 250μm D. Box plot showing quantification of weight 8 weeks post-injection of p53^+/+^ (WT) injected with GFP or PR_50_-GFP and p53^-/-^ mice injected with PR_50_-GFP (mean ± s.e.m., number of animals = 4, Student’s t-test, non-parametric, p = n.s.). E. Graph bar showing quantification of grip strength (5 measurements/animal) performed at 8 weeks post-injection of p53^+/+^ (WT) injected with GFP or PR_50_-GFP and p53^-/-^ mice injected with PR_50_-GFP (mean ± s.e.m., number of animals = 4, Student’s t-test, non-parametric, * p = 0.0274, * p = 0.0133)

**Table 2:** Mantel-Cox test summary for neuronal survival data of PR50 vs PR50-NES relative to Figure 4

| PR50 vs PR50-NES |  |  |
| --- | --- | --- |
| Log-rank (Mantel-Cox) test |  |  |
| Chi square | 76.82 |  |
| df | 1 |  |
| P value | <0.0001 |  |
| P value summary | **** |  |
| Are the survival curves sig different? | Yes |  |
| Hazard Ratio (Mantel-Haenszel) | PR50 vs PR50-NES | PR50-NES vs PR50 |
| Ratio (and its reciprocal) | 3.187 | 0.3137 |
| 95% CI of ratio | 2.460 to 4.131 | 0.2421 to 0.4066 |

We then sought to verify whether p53 is also a critical protein that mediates poly (PR) toxicity *in vivo* by taking a second approach in which we injected wild-type and homozygous p53 knockout (p53^-/-^) mice with lentivirus expressing GFP or PR_50_-GFP under an eIF2a promoter (which preferentially drives neuronal expression). The viral constructs were administered directly into the spinal cord through laminectomy and bilateral intraspinal injections in C4-C5-C6. The region targeted is the ventral horn of the C4-C6 spinal cord section, which innervates both the muscle of the forelimbs and part of the diaphragm (Lepore, 2011). Through light-sheet microscopy on cleared whole tissue, we could visualize the 6 injection sites in the spinal cord (**EV4 A**). From the dorsal view, the six injection sites are recognizable. From the latter perspective, instead, we can appreciate the depth of the injection from the dorsal horn to the ventral horn, which is the target site where motor neurons are (**Movie 1**). Mice underwent an assessment of forelimbs performance through grip strength 10 weeks after surgery. Then the spinal cord was dissected and kept for further analysis (**Fig. 5B**). We chose this time point because it is before cancer development in the p53-null mice. As expected, histology performed on tissues eight weeks post-injection showed viral expression at the injection sites in C4-C5-C6 (**Fig. 5C**).

At week 8, we recorded a small but significant (2.4 gr. on average) drop in the weight of the PR_50_-GFP injected WT mice compared to GFP injected WT mice (**Fig. 5D**). PR_50_-GFP injected mice showed lesser forelimb strength compared to GFP injected mice (**Fig. 5E**). Intriguingly, we observed increased force in the p53^-/-^ group injected with PR_50_-GFP to levels comparable to WT controls, supporting the notion that keeping p53 levels low helps prevent motor function degeneration.

Mdm2 is a ubiquitinating enzyme that maintains low levels of p53, preventing cells from initiating p53-dependent apoptosis (Marine & Lozano, 2010). We performed the intraspinal injection described above in mice overexpressing Mdm2. Overexpression of Mdm2 (Mdm2 OE) ensures that even if p53 transcription and translation are increased, p53 is ubiquitinated and targeted for proteasomal degradation, thereby preventing activation of downstream apoptotic pathways (Jones *et al*, 1998). Mdm2 OE mice develop tumors much later in life than p53^-/-^ mice (Jones *et al*., 1998), enabling behavioral assessment at later time points. We measured forelimb gaiting by DigiGait analysis (**EV5 A**).

Interestingly, degeneration in Mdm2 OE mice was delayed between weeks 6 and 9 (**EV5 B-E**), as shown by the differences in gaiting parameters such as propel, stance, stride, and stride frequency. Also, immunofluorescence performed on longitudinal sections clearly show that around the three sites of injections, there is pronounced neuronal loss as detected by loss of NeuN in both Mdm2 OE and WT animals at the endpoint (**EV4 B**).

## DISCUSSION

In this study, we demonstrated the central role of nucleolar stress and the critical role of p53 in PR-mediated neurotoxicity. The robust neurotoxic phenotype of PR could be mitigated by manipulating PR nuclear localization or suppressing the pathways downstream of the PR-mediated nucleolar stress. It is known that nucleolar abnormalities are common in other neurodegenerative diseases such as Parkinson’s disease (Parlato & Liss, 2014). Nucleolar stress is a feature of ALS/FTD associated with C9orf72 and some sporadic cases (Mizielinska *et al*, 2017). Nucleolar abnormalities have not been shown in ALS cases caused by SOD1, TDP-43, FUS, and NEK-1 mutations (Konopka & Atkin, 2022), although significant DNA damage, a type of cellular insult known to promote nucleolar stress and to activate the p53 pathway (Bordoni *et al*, 2019; Higelin *et al*, 2018; Mitra *et al*, 2019; Wang & Hegde, 2019), was reported. The arginine-rich DPRs, PR, and GR tend to localize in membrane-less structures like the nucleolus (Hartmann *et al*, 2018; Konopka & Atkin, 2022; Lee *et al*., 2016) as these structures are rich in proteins containing low complexity domains and the interaction of de-structured peptides like GR or PR with these proteins is favored (Boeynaems *et al*, 2017). Arginine-rich DPRs expressing transgenic mouse models displayed nucleolar localization of these dipeptides in neurons. This is particularly evident in the case of PR (Zhang *et al*., 2019; Zhang *et al*., 2018). The nucleolar localization pattern was also recapitulated in cell and fly models (Mizielinska *et al*., 2017). In our study, PR did not affect nucleoli’s morphology and size, whereas it changes the liquid droplet-like properties when expressed in rat cortical neurons. This evidence validates, giving disease relevance to what has been published thus far for this DPR (Hartmann *et al*., 2018; Lee *et al*., 2016; Zhang *et al*., 2019) in non-neuronal cell lines. While post-mortem tissue from C9-ALS patients did not show evidence for GR and PR in the nucleolus of the still identifiable neurons (Mizielinska *et al*., 2017), there was clear evidence for nucleolar changes in volume, suggesting a direct involvement of the nucleolus is occurring in the disease. Irrespective of the pathology of the nucleolus, the evidence points to nucleolar stress as one of the critical players of C9-ALS neurodegeneration (Herrmann & Parlato, 2018). Therefore, it is essential to investigate the pathways downstream of nucleolar stress to discover potential therapeutic targets. Thus far, senescence is one of the most understudied molecular pathways associated with nucleolar stress. Senescence is an intermediate status common among replicative cells and characterized by a shift in cell metabolism, cell growth arrest, and production of a specific secretome (cytokines and extracellular vesicles) (González-Gualda *et al*, 2021; Martínez-Cué & Rueda, 2020; Saez-Atienzar & Masliah, 2020). This state is caused by multiple stimuli, such as ROS production, DNA damage, and protein aggregates, all sensed by the nucleolus (Van Deursen, 2014). Many reports show that neurons also undergo a state that resembles senescence (Casadesus *et al*, 2012; Jurk *et al*., 2012; Ohashi *et al*, 2018; Shaker *et al*., 2021). Senescence has been indeed associated with the pathogenesis of many neurodegenerative diseases, such as (Parkinson’s disease (PD) (Ho *et al*, 2021) and Alzheimer’s disease (AD) (Wei *et al*, 2016). Evidence of senescence was also detected in ALS post-mortem tissues (Vazquez-Villasenor *et al*., 2020) and the human mutant SOD1^G93A^ transgenic mouse model of ALS(Trias *et al*, 2019).

Interestingly, clearance of senescent astrocytes in the central nervous system slowed the progression of cognitive decline in AD (Bussian *et al*, 2018). Here, we did not detect activation of the senescent pathway in neurons by PR alone. Indeed, even if we measured a robust increase in p53 levels, this did not result in measurable senescence. As senescence is not only triggered by p53, we also evaluated p16- and NF-kB-associated pathways, finding no difference between PR-expressing neurons and controls.

As C9-ALS/FTD is a multifactorial disease (Gendron & Petrucelli, 2017), other pathological events might be acting in concert to trigger the senescence phenotype in neurons (e.g., the presence of other DPRs, C9orf72 haploinsufficiency, RNA foci). It would be interesting to pursue the study of senescence in models which encompass all the C9-ALS/FTD features and undergo longer, less artificial in-vitro aging, such as animal models or cerebral organoids. In the transgenic mutant SOD1 mouse model of ALS, the onset of a senescence-like phenotype has been reported for non-neuronal cells (Trias *et al*., 2019).

Persistent nucleolar stress and increased p53 levels could also lead to apoptosis. Apoptosis has been demonstrated to be the critical pathway of PR-associated toxicity by us and others (Maor-Nof *et al*, 2021). Intervening both upstream, as by genetically increasing Mdm2-dependent ubiquitination and degradation of p53, or downstream, as presented by others, leads to reducing neuron toxicity associated with PR. After confirming that p53 knock-out can partially rescue the PR-associated motor behavioral phenotype, we harnessed a p53 regulatory pathway, specifically Mdm2, to indirectly affect p53 levels and activity and rescue disease phenotypes. Mdm2 is a ubiquitinase that targets p53 for proteasomal degradation, keeping p53 levels in check (Marine & Lozano, 2010). When the nucleolus is under stress, two main events happen that cause the release of Mdm2 from p53, causing increased p53 levels. The first is the release of free ribosomal proteins such as Rpl11, which bind Mdm2 (Sasaki *et al*, 2011), and the second is the release of NPM1, which also binds Mdm2 (Box *et al*, 2016). Here we show that overexpression of Mdm2, using a transgenic mouse model, also slows motor impairment caused by PR. Contrary to the ablation of p53 (Jacks *et al*, 1994), Mdm2 overexpression causes a much slower development of neoplasia (Jones *et al*., 1998), making local and transient stabilization a potentially more suitable therapeutic target.

Overall, our study has offered the rationale to implement future therapeutic strategies to counteract the robust neurotoxicity of C9orf72-ALS/FTD-linked arginine-rich DPRs like PR at three different levels: screening for drugs that rescue nucleolar stress; drugs that prevent PR localization to the nucleolus; lastly, drugs that rescue apoptosis and promote Mdm2 activity and p53 degradation in a CNS specific manner.

## MATERIALS AND METHODS

### Plasmids and transfection

The following plasmids were used: pcDNA3_FLAG_GFP(Wen *et al*., 2014), pcDNA3_FLAG_PR50_GFP, pcDNA3_FLAG_3XNES_PR50_GFP, GFP-NPM WT (Addegene #17578), pcDNA3_hSyn_Td-Tomato^(Bonizec *et al*, 2014)^, pcDNA3_FLAG_GP50_GFP(Bonizec *et al*., 2014), pCDNA3 (Addgene plasmid # 128034), T7-p53DD-pcDNA3, referred to here as DN-p53, was a gift from William Kaelin (Addgene plasmid # 25989; http://n2t.net/addgene:25989; RRID:Addgene_25989). To generate pcDNA3_FLAG_3XNES_PR50_GFP, a fragment containing the three repetitions of the NES was synthesized by Azenta into a plasmid backbone. This region was flanked by BamHI and EcoRI, which were used to clone the sequence in the pcDNA3_FLAG_PR50_GFP between PR50 and the sequence for GFP.

pcDNA3_FLAG_mCherry, pcDNA3_FLAG_PR_50__mCherry were obtained by cloning the mCherry sequence in pcDNA3_FLAG_GFP, pcDNA3_FLAG_PR_50__GFP using BamHI and XbaI flanking the GFP sequence.

Transfection was performed according to previously published protocols(Wen *et al*., 2014). Briefly, 2 μL of Lipofectamine 2000 (Invitrogen™ #11668027) were mixed with 1 μg of DNA plasmids mix diluted in OptiMEM (Gibco™ #11058021) added to the neuronal culture, incubated for 1 h. at 37°C), the medium was replaced with a fresh one. 400 ng of DNA/150,000 cells were used for all the DPR-expressing plasmids and controls. 400ng/150,000 cells were used for GFP-NPM WT. 200ng/150,000 cells were used for Td-tomato. 400ng/150,000 cells were used for p53_DN.

### Virus production and cells transduction

All viruses were produced using HEK293T culture in DMEM high glucose (Gibco™ #11965092 supplemented with 10% fetal bovine serum (Gibco™ #10082147), Penicillin/Streptomycin solution (Gibco™ #15140122) and glutamine (Gibco™ #25030081). 4ug of pLenti_hSyn_FLAG_GFP or pLenti_hSyn_FLAG_PR_50__GFP were mixed with 8ug of psPAX2 (Addgene Plasmid #12260) and 16ug of pMD2.G (Addgene Plasmid #12259) in 1mL of OptiMEM. 80uL of PEIMAX (Polysciences #49553-93-7) were then added, and the mixture was added to a 150mm culture dish with 70% confluent HEK293T. 48 hours after transfection, media was harvested and centrifuged for 5 minutes at 500 rcf to remove cells and debris. To concentrate the virus, Lentivirus Concentrator (OriGene technologies Cat # TR30025) was added to the supernatant, and the mixture was left overnight at 4°C. the mixture was then centrifuged, and the pellet was resuspended in 1:100^th^ of the initial volume, aliquoted and stored at −80°C.

Viral particles were thawed on ice before use, and 1ul/150,000 cells were added to the neuronal culture. After 48 hours, cells were harvested or fixed.

### Primary cortical neuron culture

Rat primary cortical neurons were prepared as follows. E16 pups were explanted from the breeder’s womb, heads were collected, and brains were dissected. After careful removal of meninges, tissues were chopped with scissors and then incubated in HBSS without calcium and magnesium (ThermoScientifìc™ #88284) with 0.2% of trypsin (Gibco™ #15090046) for 45 minutes at 37°C on an orbital shaker (low rcf to avoids stressing the culture). FBS was added to stop trypsin activity, and cells were then homogenized by pipetting and centrifuged at 800rpm for 10 minutes at 4°C. Cells were rewashed with 10 mL of HBSS and then strained through a 70μm filter. Cells were counted and plated: 12-well plate with 300,000 cells/well, 24-well plate with 150,000 cells/well, 35mm dish 35,000/dish. Neurons were cultured at 37°C, 5% CO_2_ in neurobasal supplemented with B27 and Penn/Strep solution. When the genotype of the culture was unknown (like in the case of p53 mice), each brain was kept separated.

### Drosophila Melanogaster Assay

Plasmids: To make the NES and NLS plasmids, cassettes were PCR amplified from the pUAST-Flag-DPR_50_-eGFP plasmids(Wen *et al*., 2014) using the forward primer MCS_of_PUAST which contains NotI site and reverse primers with homology to the 3’ end of eGFP and an NES and an XbaI site (called GFP-NES). This cassette was then inserted into pUAST-Flag-RAN50-eGFP digested with NotI and XbaI. The consensus NES was created with help from NES base (LTWRLNRLIL).

D. melanogaster maintenance and genetics: All Drosophila stocks were cultured and maintained on standard dextrose media on a 12-hour light/dark cycle at a temperature of 18°C until otherwise noted. Transgenic lines used in this study were: pUAST attb FLAG eGFP, attb UAS FLAG PR_50_-eGFP 86F8, pUASTattb FLAG eGFP NES, pUASTattb FLAG PR_50_ eGFP NES. The GMR GAL4 driver line was obtained from Bloomington Drosophila Stock Center. Transgenes and markers were inserted at defined locations in the genome using phiC31 mediated transgenesis by the company Bestgene Inc as previously reported(Casci *et al*, 2019; Kour *et al*, 2021; Piol *et al*, 2023; Ramesh *et al*, 2020a; Ramesh *et al*, 2020b).

Eye Degeneration: Expression of the transgenes in the Drosophila eye was driven using the GMR Gal4 driver at 18°C. One day old adults were collected, anesthetized and external images of the eye were taken using LEICA M205C microscope equipped with a LEICA DFC450 camera. The eye phenotype of > 20 animals per genotype was scored in an unblinded manner (Casci *et al*., 2019; Fortuna *et al*, 2021; Piol *et al*., 2023; Ramesh *et al*., 2020a; Ramesh *et al*., 2020b) and the differences between the genotypes were assessed using One Way ANOVA followed by Tukey’s multiple comparison test.

### Animals

SAS Sprague Dawley Rats from Charles River were used to obtain rat primary cortical neurons. They were housed according to the IACUC guidelines and approved protocol. Dr. Eischen kindly provided B6.129S2-Trp53tm1Tyj/J and C57Bl/6 p53^-/-^ mice previously reported and obtained from Dr. Guillermina Lozano. According to published protocols, genotyping of these animals was performed 14-16 days after birth.

C57Bl/6 Mdm2 transgenic mice were previously reported and provided by Dr. Steven Jones(Jones *et al*., 1998). Wild-type littermate controls were used. Animals were genotyped according to published protocol(Jones *et al*., 1998)Animals were housed and bred according to the IACUC guidelines and approved protocol.

### Intraspinal injection

The Thomas Jefferson University IACUC committee revised and approved the following procedure. Animals were anesthetized with 2% isoflurane and kept under anesthesia during the entire surgery. The skin was cut open with a razor blade to expose the muscles covering the spinal columns. Muscles were then carefully taken apart to reveal the column. C4, C5, and C6 vertebrae were opened to reveal the spinal cord. 2 ul of virus (pLenti_hSyn_FLAG_GFP or pLenti_hSyn_FLAG_PR_50__GFP) were injected in the ventral horn with a 33-gauge needle on a microsyringe controlled by an UltraMicroPump and Micro4 Microsyringe Pump Controller (World Precision Instruments, Sarasota, FL, USA). A total of six injections were performed bilaterally from C4 to C6. After injection, muscles were sutured, and skin was clipped following standard procedure.

### Behavioral tests

Grip strength: animal forelimbs were placed on a bar, and tension was applied by pulling the mouse’s tail. Resistance right before paws detach from the bar was registered.

Five measurements per mouse were taken each time.

Hanging wire: the animal was placed on a grid which was then flipped, leaving the animal upside down. Animals were left tested for two minutes total. The time before the animal detached from the grid was recorded, and three measurements per mouse were recorded.

Gaiting analysis: DigiGait apparatus was used. Animals were placed on a treadmill and recorded for 5 sec. Each movie was then analyzed with DigiGait software.

### Immunofluorescence staining

Cells were fixed for 20 minutes at 37° with 4% PFA. After a wash with PBS, cells were permeabilized and blocked in one step using PermBlock solution (1% FBS, 1% BSA, 0.05% Triton X100) for 1 hour at room temperature. Antibodies were diluted 0.1% of BSA and incubated with cells overnight at 4° C. After two washes in PBS, cells were exposed to secondary antibodies in BSA 1%. After two final washes in PBS, cells were treated with Hoechst (Abcam #ab228550) to stain the nuclei (1:3000 in PBS).

Perfused tissues were post-fixed overnight at 4° C in 4% PFA. They were cryoprotected with 30% sucrose solution and then embedded in OCT. After sectioning, 40μm free floating slices were incubated with 0.05% tritonX100 in PBS for 1 hour at room temperature. Slices were then incubated with 1% goat serum and 0.05% Triton X100 in PBS for 2 hours at room temperature. Antibodies were diluted in antibody solution (0.025% of FBS, 0.015% of Triton X100 in PBS) and incubated with the slices overnight at 4° C. After two washes with PBS, slices were incubated with secondary antibodies diluted in antibody solution. Slices were then rinsed in PBS twice and incubated with Hoechst solution (1:3000 in PBS). Slices were then mounted on glass slides with AquaMount (VWR #41799-008) for acquisition.

### Beta-galactosidase staining, acquisition, and analysis

Fixed cells were stained following previously published protocols. Fixed cells were incubated with staining solution (1mg/mL of X-Gal,5-Bromo-4-chloro-3-indolyl-beta-D-galactopyranoside; Sigma Aldrich #11680293001)(Lorenzo, 2013) for 16 hours at 37°. Cells were washed with PBS, and images were acquired with EVOS (Thermo Fisher). 3 fields/well images were obtained, and the number of cells positive for β-galactosidase per image was counted and graphed.

### Confocal imaging and analysis

Images were acquired using a confocal Nikon A1R microscope. 10 nuclei/condition were imaged for each experiment. ROIs were built around nucleoli or nuclei, and NIS Nikon software calculated brightness and dimension.

### FRAP

10 cells/experiment double positive for GFP-NPM and mCherry were acquired. Images were taken with a 60X objective with a 3X digital zoom. ROI was drawn around a single nucleolus, and after 5 seconds of baseline recording, a 100% power blue laser light was applied to photobleach GFP fluorescence. Images were recorded for 90 sec after photobleaching. Data normalization, taking background and whole cell area, was performed by FRAPeasy. Data were then plotted using Prism GraphPad 9.0. statistical analysis was performed by best-fitting the normalized mean fluorescence against time and by non-parametric t-test between the calculated slopes of the two curves.

### Live imaging

Neurons double positive for green and red fluorescence were imaged 24 hours after transfection and then every 24 hours with an Eclipse Ti Nikon Microscope. Neurons were tracked over time and were censored as dead upon dendrite and cell body fragmentation. Kaplan Meier curves were built using Prism 9.0.

### qPCR

RNA isolation was performed using Trizol (Ambion #15596026) according to the manufacturer’s instructions. We used Nanodrop to measure RNA concentration. 1ug of total RNA was treated with DNaseI (Invitrogen™ #18068015) and then retrotranscribed using RTIII (Invitrogen™ #18080044). 25ng of cDNA were used for SYBRgreen (Applied Biosystems™ #4309155) qPCR. Used primers are in Table 3.

**Table 3:**
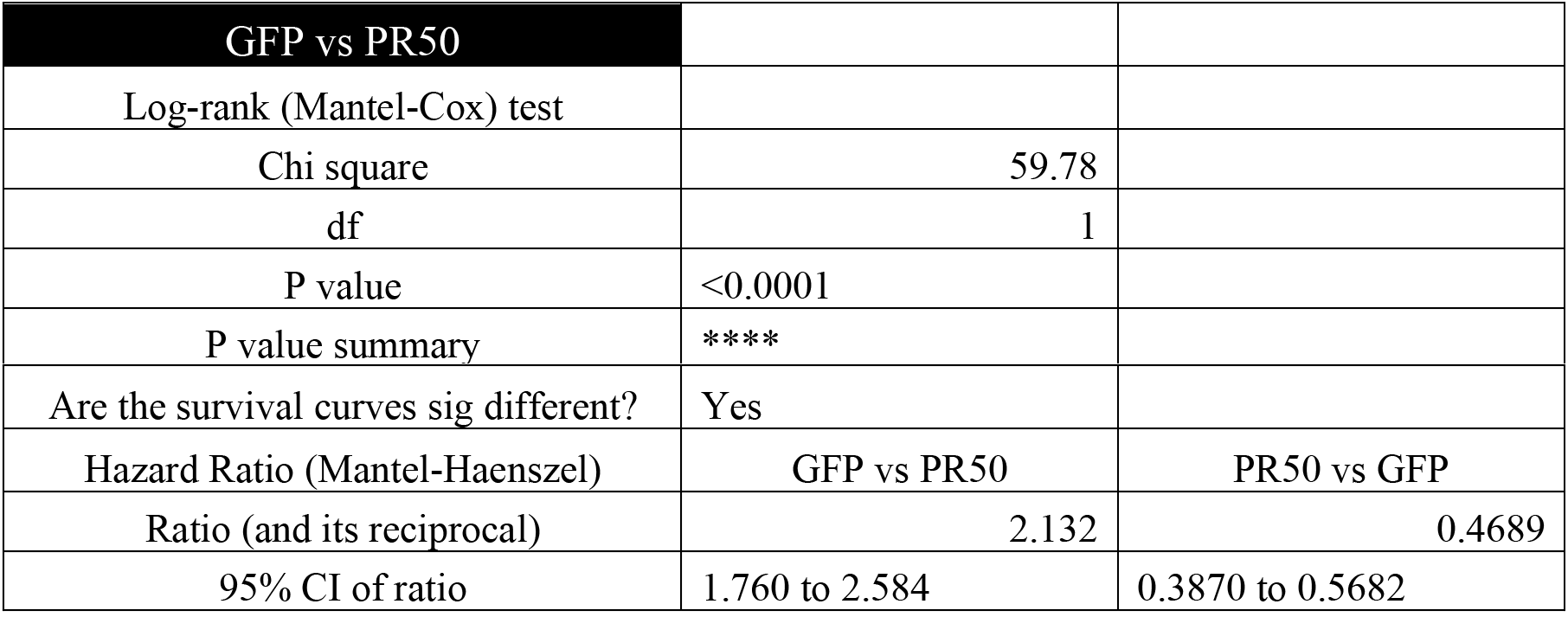
Mantel-Cox test summary for neuronal survival data of GFP vs PR50 relative to Figure 5

| GFP vs PR50 |  |
| --- | --- |
| Log-rank (Mantel-Cox) test |  |
| Chi square | 59.78 |
| df | 1 |
| P value | <0.0001 |
| P value summary | **** |

|  |  |  |
| --- | --- | --- |
| Are the survival curves sig different? | Yes |  |
| Hazard Ratio (Mantel-Haenszel) | GFP vs PR50 | PR50 vs GFP |
| Ratio (and its reciprocal) | 2.132 | 0.4689 |
| 95% CI of ratio | 1.760 to 2.584 | 0.3870 to 0.5682 |

**Table 4:** Mantel-Cox test summary for neuronal survival data of PR50 vs PR50/p53DN relative to Figure 5

| PR50 vs PR50/DN-p53 |  |  |
| --- | --- | --- |
| Log-rank (Mantel-Cox) test |  |  |
| Chi square | 35.12 |  |
| df | 1 |  |
| P value | <0.0001 |  |
| P value summary | **** |  |
| Are the survival curves sig different? | Yes |  |
| Hazard Ratio (Mantel-Haenszel) | PR50 vs PR50/DN-p53 | PR50/DN-p53 vs PR50 |
| Ratio (and its reciprocal) | 1.754 | 0.57 |
| 95% CI of ratio | 1.457 to 2.113 | 0.4733 to 0.6864 |

**Table 5:**
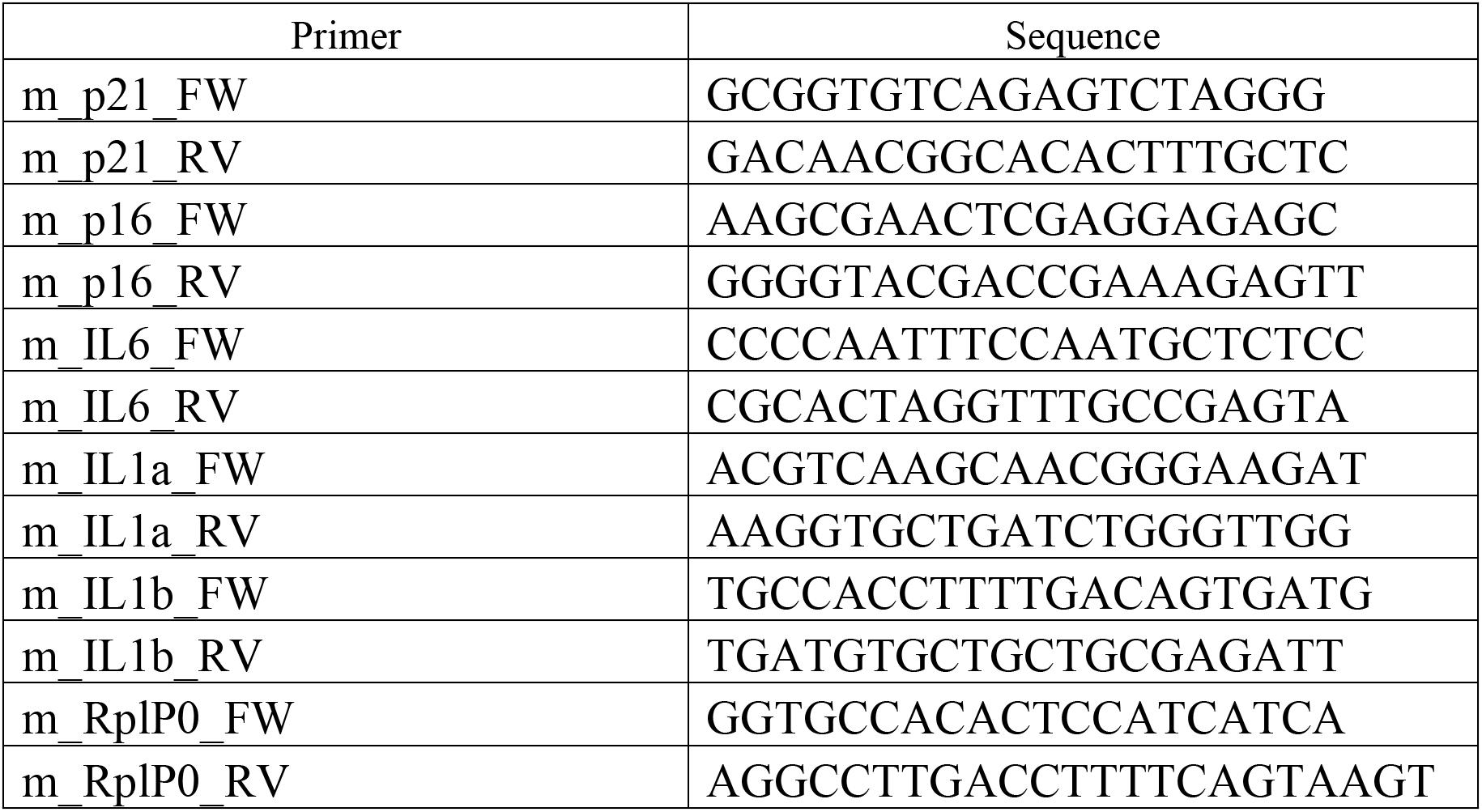
sequences of the used primers

### ELISA

Colorimetric-based ELISA (R&D Systems #M6000B) kit was used to measure IL-6 concentration in cortical neuron media. Media was collected and centrifuged at 500 rcf for 10 minutes at 4°C to remove cells and significant debris. 100uL of media was used for ELISA. The experiment was carried out following manufacturer instructions, and IL6 concentration was interpolated using a concentration curve.

### Western blot

Cells were harvested and lysed in RIPA buffer. After 15 minutes on ice, samples were sonicated and centrifuged at 16,000 rpm for 10 minutes at 4°C. Protein concentration was measured using the BCA assay. 15ug of total proteins/sample was loaded in a precast 4-15% polyacrylamide gel (Bio-Rad #4561033EDU) and run at 100mV for 1 hour. Before transferring, gels were activated using UV to obtain total protein staining. Gels were then transferred on nitrocellulose filter paper (0.45μm pores; Bio-Rad #1620115) using a Trans-Blot Turbo Transfer System apparatus (Bio-Rad). Membranes were then imaged again for total protein and blocked in 5% milk solution in TBS-T 1X for 1 hour at room temperature. Membranes were then incubated o/n at 4°C with primary antibodies diluted in 5% BSA solution in TBS-T 1X. After two washes in TBS-T 1X at room temperature, membranes were incubated with a secondary antibody (1:10,000). Membranes were washed four times and then exposed to ECL solution (Thermo Fisher Scientific #34096). Images were acquired using ChemiDoc XRS+ System (Bio-Rad). The Image Lab software was used for WB quantification to quantify the band intensity for the protein of interest. Relative quantification was performed using total protein staining or loading control protein such as GAPDH/Actin. WB data were then processed to show a fold change compared to GFP.

### Whole tissue clearing and imaging

Whole tissue staining and clearing on spinal cords were performed following a previously published protocol(Chi *et al*, 2018). After perfusion and dissection, spinal cords were fixed overnight and then washed in PBS for 24 hrs. The tissues were then passed through multiple ethanol gradients from 20% to 80% to dehydrate them. Using 4% bleach in acetic acid, the tissue was then bleached for 4 hours, rehydrated, and permeabilized with 0.02% Tween 20 for one day at 37°. Primary antibody (NeuN) was incubated for three days under constant shaking at 37°. After 4 washes of two hours each, samples were incubated with the secondary antibody for three days under continuous shaking at 37°. After 4 washes of two hours each, samples were dehydrated with increasing methanol concentration and delipidated with DCM. The tissue was then put in DBE for clearing and, the day after was imaged with a UM Blaze (Miltenyi). Images were first acquired with a 4X objective with 0.6X zoom. To achieve higher magnification, images were then taken with a 12X objective. Image assembling and rendering were performed using Imaris.

### Antibodies

P53 (WB dilution 1:1000; IF dilution 1:500 Cell Signalling Technology #2524S), p21 (WB dilution 1:500 Cell Signalling Technology #2947S), p16 (WB dilution 1:500 Santa Cruz Biotechnology #S1661), NfkB (WB dilution 1:1,000 Abcam #16502), B23/NPM (IF dilution 1:2,000 Sigma-Aldrich #B0556), Map2 (IF dilution 1:2,000 Novus Biologicals #NB300-213), GAPDH (WB dilution 1:5000 Fitzgerald #10R-G109A), GFP (WB dilution 1:1,000 Proteintech #50430-2-AP), NeuN (IF dilution 1:2,000 Novus Biologicals #NBP1-92693), anti-rabbit HRP (WB dilution 1:10,000 Cytiva #NA9340), anti-mouse HRP (WB dilution 1:10,000 Cytiva #NA9310), anti-mouse 546 Alexa Fluor (IF dilution 1:1,000 Life Technologies #A10036), antichicken 647 Alexa Fluor (IF dilution 1:1,000 Life Technologies #A-21463)

### Statistical analysis

All the experiments were carried out with three or more biological replicates (*n*). More than 10 technical replicates (*m*) per *n* were included in the analysis. If two groups were compared, a student *t*-test was used, and one-way ANOVA was used when more than two groups were compared.

## AUTHOR CONTRIBUTION

MEC and DT planned the experiment and interpreted the data. MEC carried out in vivo and in vitro experiments, performed data analysis, and wrote the manuscript. JH carried out the studies with p53-DN and qPCR for pre-processed rRNA. MD performed the experiments in the flies. KK conducted the nuclear p53 quantification study. SSM prepared primary cortical neurons. AB discussed and suggested an experiment on senescence and revised the manuscript. ATN designed and produced lentivirus. LTG performed and imaged senescence-associated beta-galactosidase assay. VK maintained the colony and performed genotyping. CME provided the mice and expertise on Mdm2 and p53 to guide the studies. UBP, CH, ARH, CME, and PP revised the manuscript. DT supervised the entire research and revised the manuscript.

## ACKNOWLEDGEMENTS

The following sources supported this work: National Institutes of Health R21-NS090912 (D.T.), RF1-AG057882 (D.T.), R01-NS109150 (P.P.); Muscular Dystrophy Association grant 628389 (D.T.); DoD grant W81XWH-21-1-0134 (M.E.C.); Herbert A. Rosenthal endowed Chair fund (C.M.E.), and Farber Family Foundation (D.T., P.P).

## EXPANDED VIEW FIGURES LEGENDS

**Expanded View 1.**
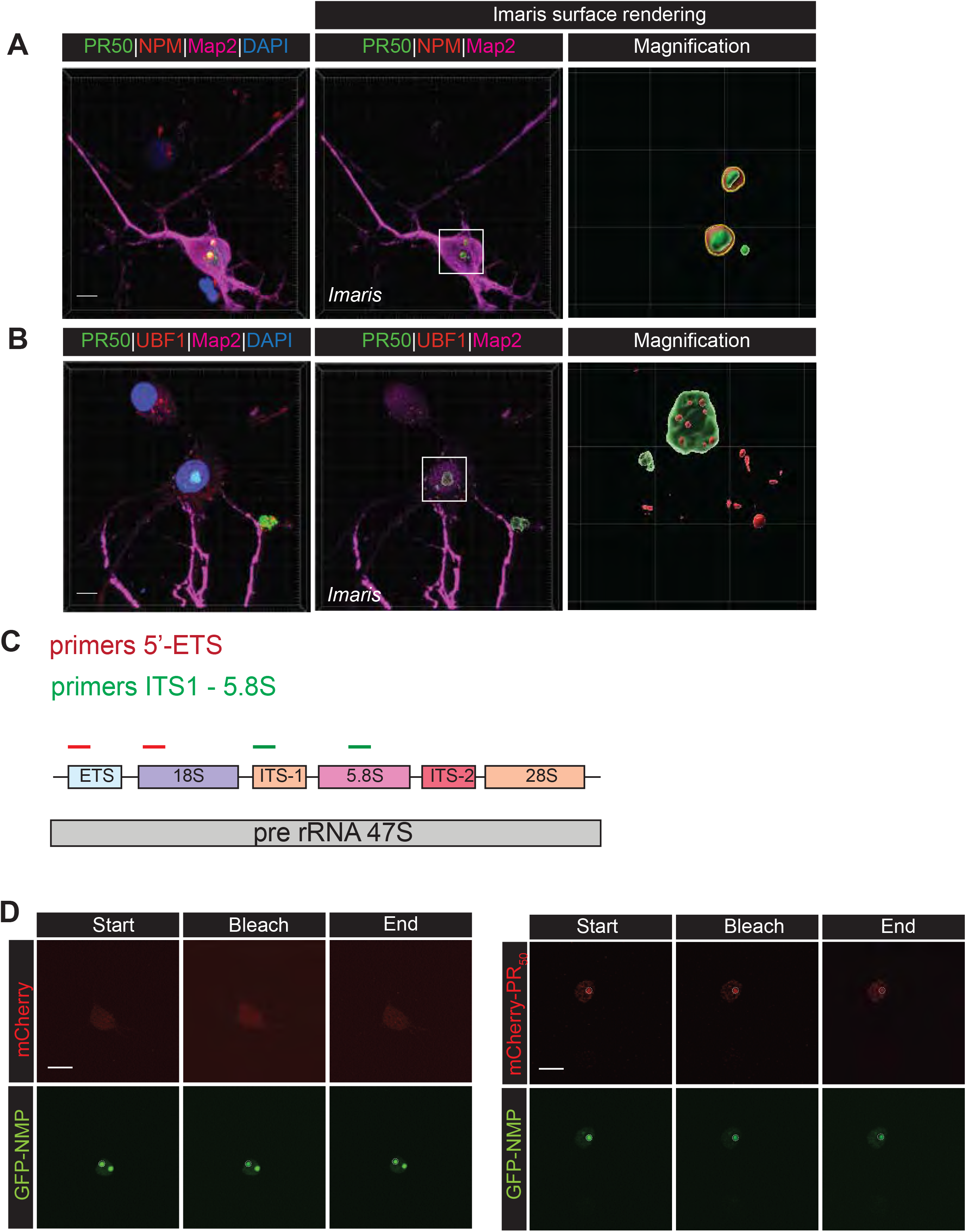
A. Confocal imaging and 3D Imaris rendering of rat primary cortical neurons transfected with PR_50_-GFP. GFP is shown in green, NPM is shown in red, Map2 is shown in magenta, and nuclei (Hoechst) are shown in blue. Scale bar = 10 μm B. Confocal imaging and Imaris rendering of rat primary cortical neurons transduced with PR_50_-GFP. GFP is shown in green, Ubf1 is shown in red, Map2 is shown in magenta, and nuclei (Hoechst) are shown in blue. Scale bar = 10 μm C. Schematic of primers used for qPCR for pre-processed rRNA. D. Representative images of nucleoli bleached for FRAP experiment (white ROI). Rat primary cortical neurons are transfected with GFP-NPM and mCherry or PR_50_-mCherry. Time points shown: start (0 sec), bleach (5 sec), and end (180 sec). Scale bar = 10μm.

**Expanded View 2.**
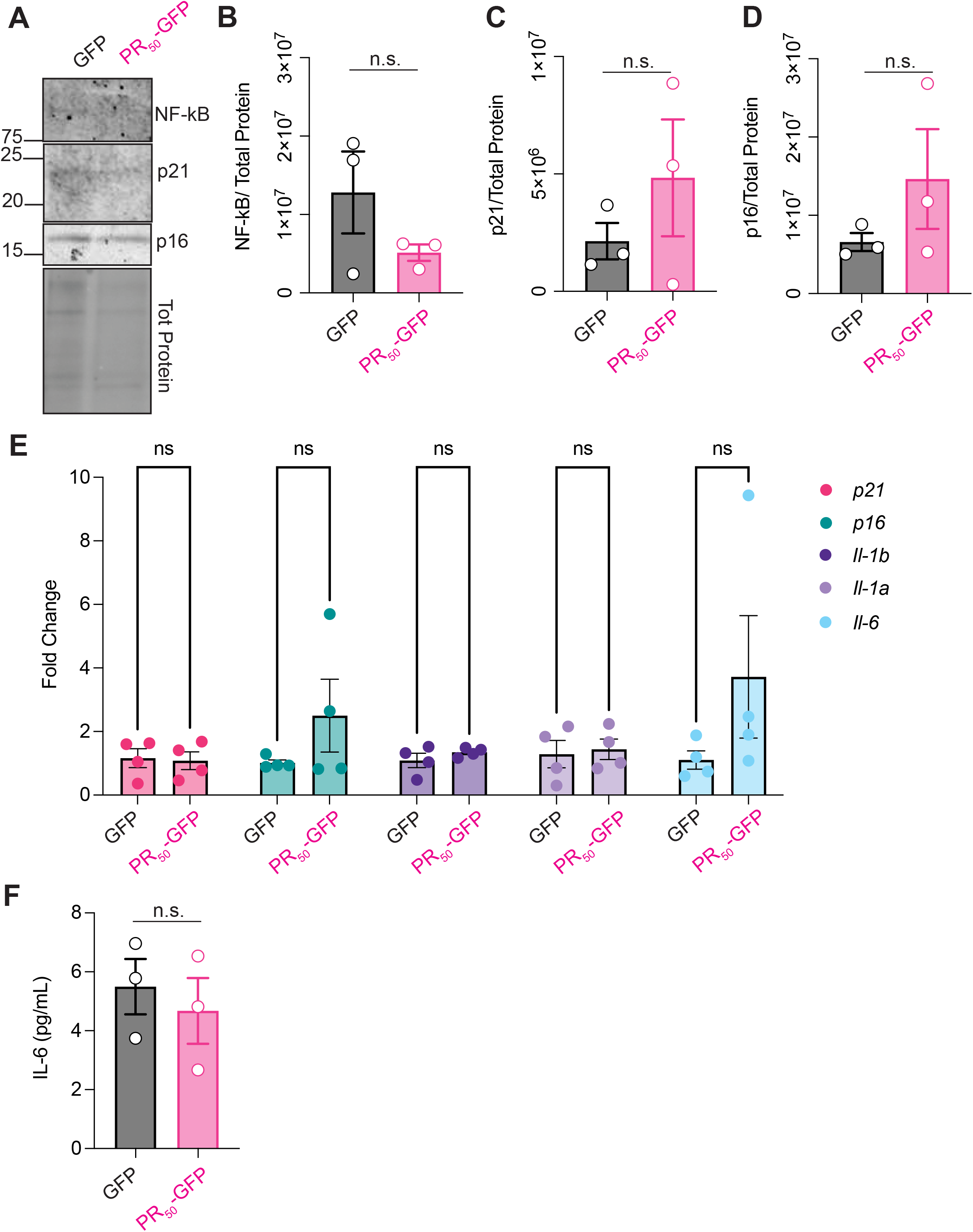
A. Western blot for p21, p16, and NF-kB and total protein performed on mouse primary cortical neurons transduced with GFP or PR_50_-GFP 7 days post-transduction. B. Graph bar showing NF-kB quantification of Western blot performed on mouse primary cortical neurons transduced with GFP or PR_50_-GFP (mean ± s.e.m., number of experiments = 3, Student’s t-test, non-parametric, p = n.s.). C. Graph bar showing p21 quantification of Western blot performed on mouse primary cortical neurons transduced with GFP or PR_50_-GFP (mean ± s.e.m., number of experiments = 3, Student’s t-test, non-parametric, p=n.s.). D. Graph bar showing p16 quantification of Western blot performed on mouse primary cortical neurons transduced with GFP or PR_50_-GFP (mean ± s.e.m., number of experiments = 3, Student’s t-test, non-parametric, p=n.s.). E. qPCR for *p21, p16, Il6, Il1a*, and *Il1b* was performed on mouse primary cortical neurons transduced with GFP or PR_50_-GFP 7 days post-transduction (mean ± s.e.m., number of experiments = 3, Student’s t-test, One-way ANOVA, p=n.s.). F. ELISA to assay Il-6 concentration in cell culture media from mouse primary cortical neurons transduced with GFP or PR_50_-GFP, 7 days post-transduction (mean ± s.e.m., number of experiments = 3, Student’s t-test, Student’s t-test, non-parametric, p=n.s.).

**Expanded View 3.**
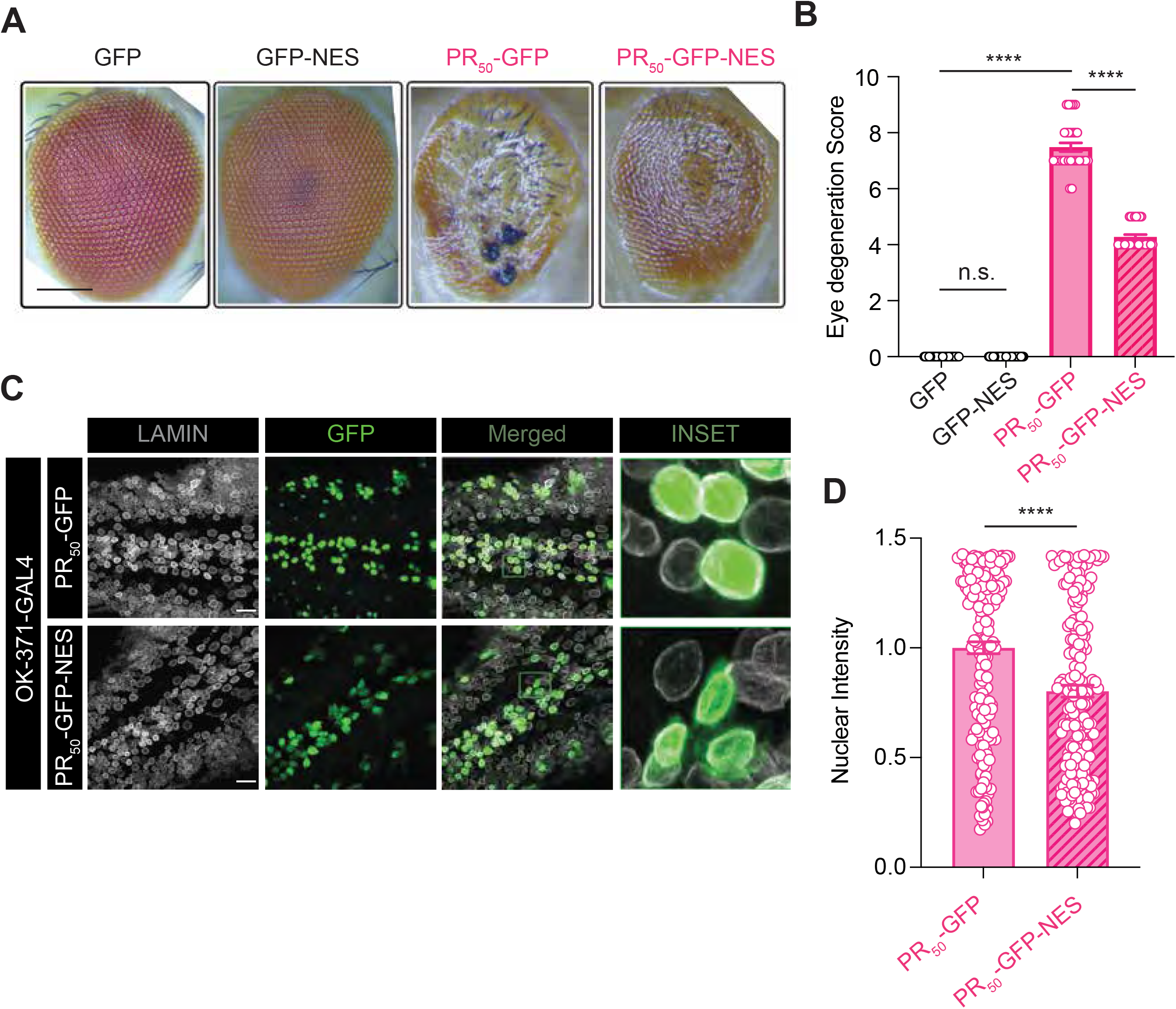
A. Stereomicroscopic images of external eyes of flies overexpressing pUAST-Flag-eGFP, pUAST-Flag-NES-eGFP, pUAST-Flag-PR_50_-eGFP, pUAST-Flag-PR_50_-NES-eGFP. Scale bar =100 μm B. Graph bar showing eye degeneration score in flies overexpressing the indicated constructs. (mean ± s.e.m., number of experiments = 3, number of flies per experiment ≥ 25, One-way ANOVA, Tukey’s multiple comparison test, **** p < 0.0001). C. Confocal images of eyes of flies overexpressing the indicated constructs stained for lamin. Lamin is shown in grey, GFP is shown in green. Scale bar=20 μm. D. Graph bar showing quantification of GFP fluorescence in flies nuclei overexpressing the indicated constructs (mean ± s.e.m., number of experiments = 1, number of flies per experiment = 20, number of cells per flies ≥ 8. Student’s t-test, non-parametric **** p< 0.0001).

**Expanded View 4.**
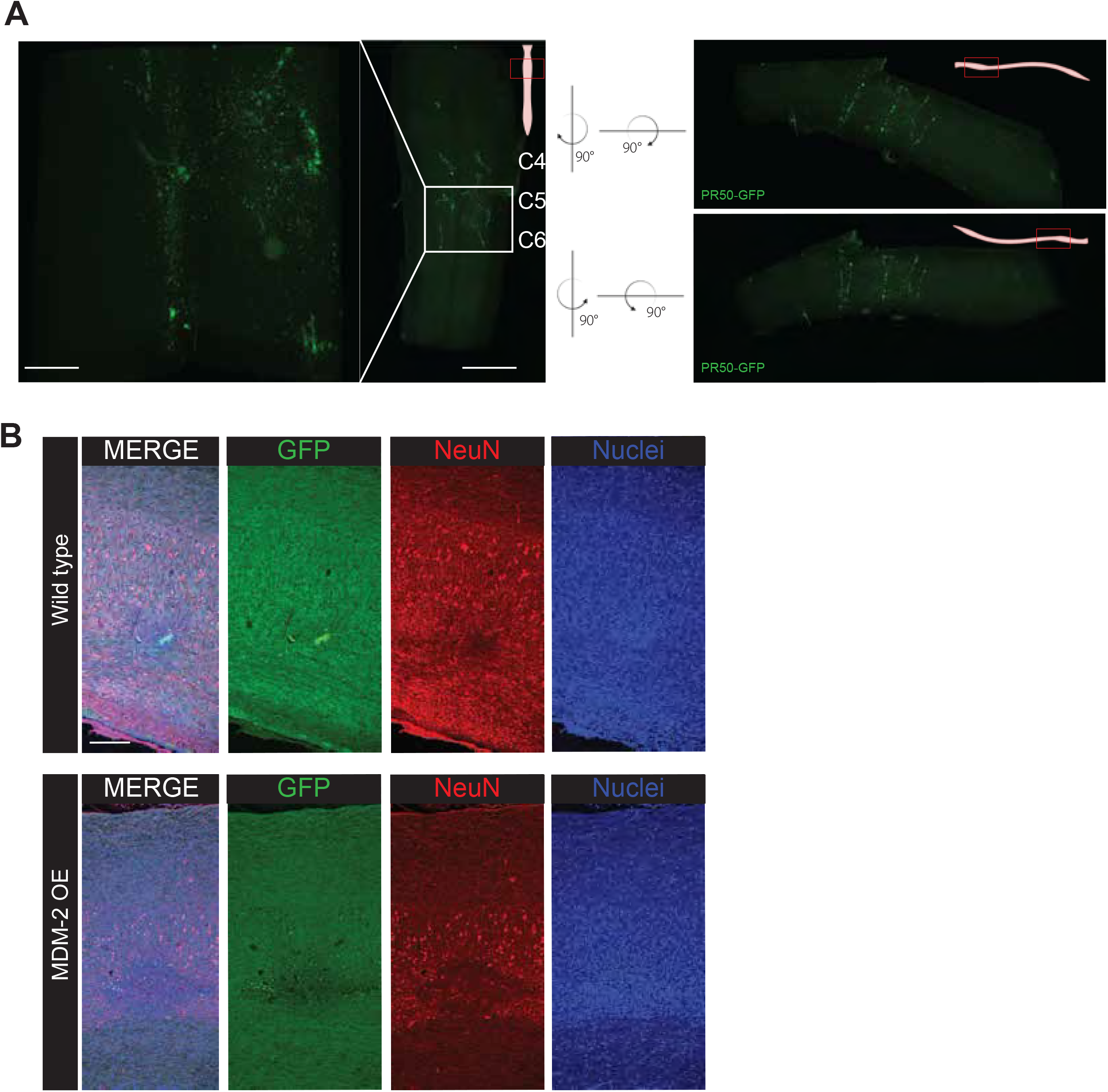
A. Center: dorsal view of spinal cord tissue C4-C6 bilateral injected with PR_50_-GFP acquired through light-sheet microscopy 4X magnification, 0.6X Zoom. Scale Bar = 0.5mm. Left: dorsal view of spinal cord tissue C4-C6 bilateral injected with PR_50_-GFP acquired through light-sheet microscopy 4X magnification, 2.5X Zoom. Scale Bar = 2mm. Right: lateral views of spinal cord tissue C4-C6 bilateral injected with PR_50_-GFP obtained through lightsheet microscopy 4X magnification, 2.5X Zoom. Scale bar = 2mm. B. Confocal imaging of PR_50_-GFP injection site in WT animal. Scale bar = 100μm C. Confocal imaging of PR_50_-GFP injection site in Mdm2 OE animal. Scale bar = 100μm

**Expanded View 5.**
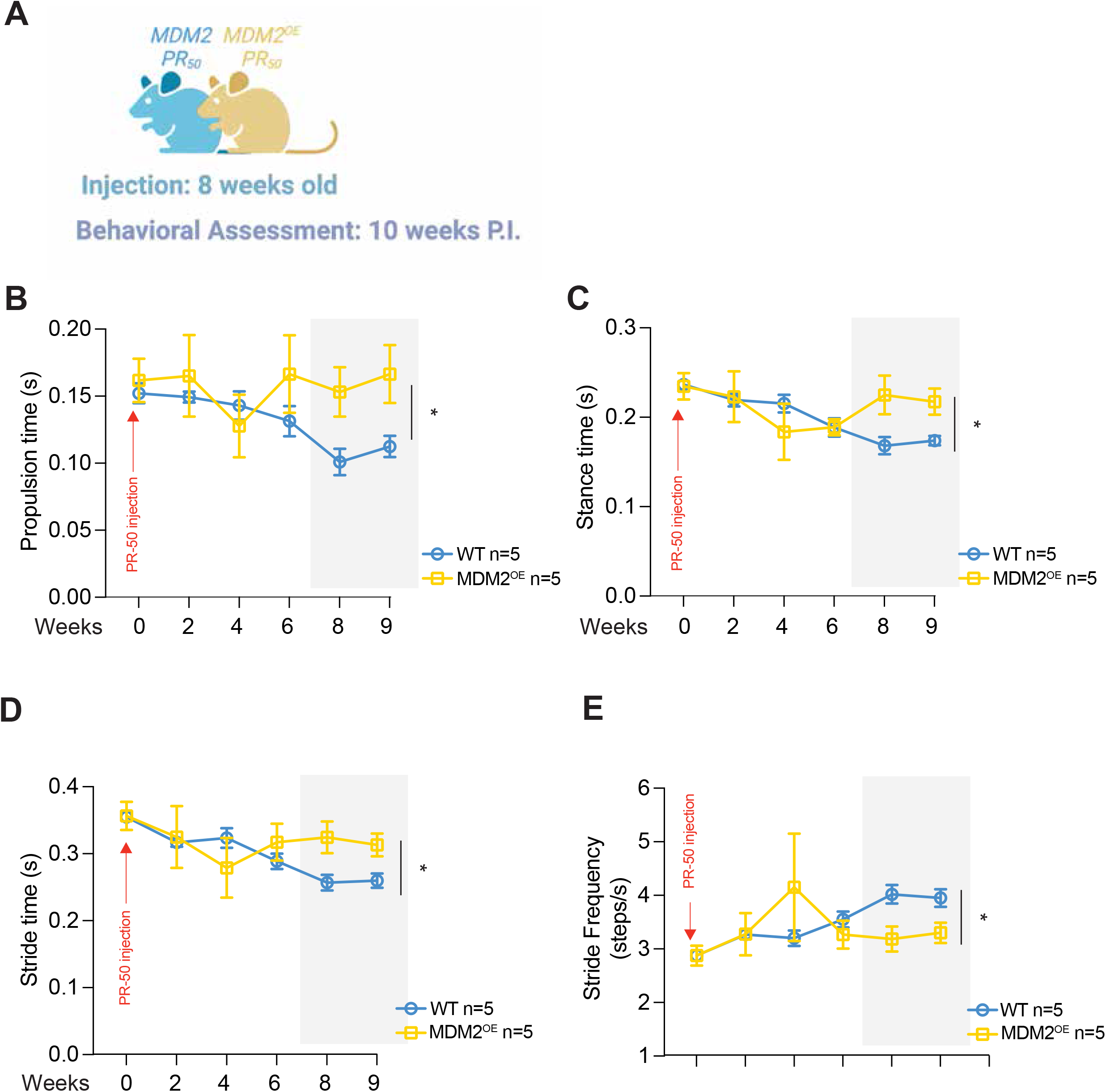
A. Schematic of the experiment. B. Over time assessment of propulsion time (s) in WT and Mdm2 OE mice injected with PR_50_-GFP. (mean ± s.e.m., number of animals ≥ 4, Student’s t-test, non-parametric, * p< 0.05). C. Over time assessment of stance time (s) in WT and Mdm2 OE mice injected with PR_50_-GFP. WT vs. Mdm2 OE p<0.05 at weeks 8 and 9 post-injection (mean ± s.e.m., number of animals ≥ 4, Student’s t-test, non-parametric, * p< 0.05). D. Over time assessment of stride time (s) in WT and Mdm2 OE mice injected with PR_50_-GFP. WT vs. Mdm2 OE p<0.05 at weeks 8 and 9 post-injection (mean ± s.e.m., number of animals ≥ 4, Student’s t-test, non-parametric, * p< 0.05). E. Over time assessment of stride frequency (steps/s) in WT and Mdm2 OE mice injected with PR_50_-GFP. WT vs. Mdm2 OE P<0.05 at weeks 8 and 9 post-injection (mean ± s.e.m., number of animals ≥ 4, Student’s t-test, non-parametric, * p< 0.05).

## REFERENCES

Ash PE, Bieniek KF, Gendron TF, Caulfield T, Lin WL, Dejesus-Hernandez M, van Blitterswijk MM, Jansen-West K, Paul 3rd JW, Rademakers R et al (2013) Unconventional translation of C9ORF72 GGGGCC expansion generates insoluble polypeptides specific to c9FTD/ALS. Neuron 77: 639–646

Biasiotto G, Zanella I (2018) The effect of C9orf72 intermediate repeat expansions in neurodegenerative and autoimmune diseases. Multiple Sclerosis and Related Disorders 27: 42–43

Boeynaems S, Bogaert E, Kovacs D, Konijnenberg A, Timmerman E, Volkov A, Guharoy M, De Decker M, Jaspers T, Ryan VH et al (2017) Phase Separation of C9orf72 Dipeptide Repeats Perturbs Stress Granule Dynamics. Molecular Cell 65: 1044–1055.e1045

Boisvert FM, van Koningsbruggen S, Navascues J, Lamond AI (2007) The multifunctional nucleolus. Nat Rev Mol Cell Biol 8: 574–585

Bonizec Ml, H??rissant L, Pokrzywa W, Geng F, Wenzel S, Howard GC, Rodriguez P, Krause S, Tansey WP, Hoppe T et al (2014) The ubiquitin-selective chaperone Cdc48/p97 associates with Ubx3 to modulate monoubiquitylation of histone H2B. Nucleic acids research 42: 10975–10986

Bordoni M, Pansarasa O, Dell’Orco M, Crippa V, Gagliardi S, Sproviero D, Bernuzzi S, Diamanti L, Ceroni M, Tedeschi G et al (2019) Nuclear Phospho-SOD1 Protects DNA from Oxidative Stress Damage in Amyotrophic Lateral Sclerosis. J Clin Med 8

Boulon S, Westman BJ, Hutten S, Boisvert FM, Lamond AI (2010) The nucleolus under stress. Mol Cell 40: 216–227

Box JK, Paquet N, Adams MN, Boucher D, Bolderson E, O’Byrne KJ, Richard DJ (2016) Nucleophosmin: from structure and function to disease development. BMC Mol Biol 17: 19

Bussian TJ, Aziz A, Meyer CF, Swenson BL, van Deursen JM, Baker DJ (2018) Clearance of senescent glial cells prevents tau-dependent pathology and cognitive decline. Nature 562: 578–582

Casadesus G, Gutierrez-Cuesta J, Lee HG, Jimenez A, Tajes M, Ortuno-Sahagun D, Camins A, Smith MA, Pallas M (2012) Neuronal cell cycle re-entry markers are altered in the senescence accelerated mouse P8 (SAMP8). J Alzheimers Dis 30: 573–583

Casci I, Krishnamurthy K, Kour S, Tripathy V, Ramesh N, Anderson EN, Marrone L, Grant RA, Oliver S, Gochenaur L et al (2019) Muscleblind acts as a modifier of FUS toxicity by modulating stress granule dynamics and SMN localization. Nat Commun 10: 5583

Chen J, Stark LA, 2019. Insights into the Relationship between Nucleolar Stress and the NF-κB Pathway, Trends in Genetics. The Author(s), pp. 768–780.

Chi J, Crane A, Wu Z, Cohen P (2018) Adipo-Clear: A Tissue Clearing Method for Three-Dimensional Imaging of Adipose Tissue. J Vis Exp

Coppé JP, Desprez PY, Krtolica A, Campisi J (2014) The Senescence-Associated Secretory Phenotype: The Dark Side of Tumor Suppression. Annual Review of Pathology: Mechanisms of Disease: 99–118

Dejesus-hernandez M, Mackenzie IR, Boeve BF, Boxer AL, Baker M, Rutherford NJ, Nicholson AM, Finch NA, Gilmer F, Adamson J et al (2011) Expanded GGGGCC hexanucleotide repeat in non-coding region of C9ORF72 causes chromosome 9p-linked frontotemporal dementia and amyotrophic lateral sclerosis. Neuron 72: 245–256

Fortuna TR, Kour S, Anderson EN, Ward C, Rajasundaram D, Donnelly CJ, Hermann A, Wyne H, Shewmaker F, Pandey UB (2021) DDX17 is involved in DNA damage repair and modifies FUS toxicity in an RGG-domain dependent manner. Acta Neuropathol 142: 515–536

Gendron TF, Petrucelli L (2017) Disease Mechanisms of C9ORF72 Repeat Expansions. 1–22

González-Gualda E, Baker AG, Fruk L, Muñoz-Espín D, 2021. A guide to assessing cellular senescence in vitro and in vivo, FEBS Journal. pp. 56–80.

Hartmann H, Hornburg D, Czuppa M, Bader J, Michaelsen M, Farny D, Arzberger T, Mann M, Meissner F, Edbauer D (2018) Proteomics and C9orf72 neuropathology identify ribosomes as poly-GR/PR interactors driving toxicity. Life Science Alliance 1: 1–13

Herrmann D, Parlato R, 2018. C9orf72-associated neurodegeneration in ALS-FTD: breaking new ground in ribosomal RNA and nucleolar dysfunction, Cell and Tissue Research. Cell and Tissue Research, pp. 351–360.

Higelin J, Catanese A, Semelink-Sedlacek LL, Oeztuerk S, Lutz AK, Bausinger J, Barbi G, Speit G, Andersen PM, Ludolph AC et al (2018) NEK1 loss-of-function mutation induces DNA damage accumulation in ALS patient-derived motoneurons. Stem Cell Res 30: 150–162

Ho DH, Nam D, Seo MK, Park SW, Seol W, Son I (2021) LRRK2 Kinase Inhibitor Rejuvenates Oxidative Stress-Induced Cellular Senescence in Neuronal Cells. Oxid Med Cell Longev 2021: 9969842

Jacks T, Remington L, Williams BO, Schmitt EM, Halachmi S, Bronson RT, Weinberg RA (1994) Tumor spectrum analysis in p53-mutant mice. Curr Biol 4: 1–7

Jones SN, Hancock AR, Vogel H, Donehower LA, Bradley A (1998) Overexpression of Mdm2 in mice reveals a p53-independent role for Mdm2 in tumorigenesis. Proc Natl Acad Sci U S A 95: 15608–15612

Jovičič A, Mertens J, Boeynaems S, Bogaert E, Chai N, Yamada SB, Paul JW, Sun S, Herdy JR, Bieri G et al (2015) Modifiers of C9orf72 dipeptide repeat toxicity connect nucleocytoplasmic transport defects to FTD/ALS. Nature Neuroscience 18: 1226–1229

Jurk D, Wang C, Miwa S, Maddick M, Korolchuk V, Tsolou A, Gonos ES, Thrasivoulou C, Jill Saffrey M, Cameron K et al (2012) Postmitotic neurons develop a p21-dependent senescence-like phenotype driven by a DNA damage response. Aging Cell 11: 996–1004

Konopka A, Atkin JD, 2022. DNA Damage, Defective DNA Repair, and Neurodegeneration in Amyotrophic Lateral Sclerosis. pp. 1–15.

Kour S, Rajan DS, Fortuna TR, Anderson EN, Ward C, Lee Y, Lee S, Shin YB, Chae JH, Choi M et al (2021) Loss of function mutations in GEMIN5 cause a neurodevelopmental disorder. Nat Commun 12: 2558

Lee K-H, Zhang P, Kim Hong J, Mitrea DM, Sarkar M, Freibaum BD, Cika J, Coughlin M, Messing J, Molliex A et al (2016) C9orf72 Dipeptide Repeats Impair the Assembly, Dynamics, and Function of Membrane-Less Organelles. Cell 167: 774–788.e717

Lee YB, Chen HJ, Peres JN, Gomez-Deza J, Attig J, Štalekar M, Troakes C, Nishimura AL, Scotter EL, Vance C et al (2013) Hexanucleotide repeats in ALS/FTD form length-dependent RNA Foci, sequester RNA binding proteins, and are neurotoxic. Cell Reports 5: 1178–1186

Lepore AC (2011) Intraspinal cell transplantation for targeting cervical ventral horn in amyotrophic lateral sclerosis and traumatic spinal cord injury. J Vis Exp

Liu F, Morderer D, Wren MC, Vettleson-Trutza SA, Wang Y, Rabichow BE, Salemi MR, Phinney BS, Oskarsson B, Dickson DW et al (2022) Proximity proteomics of C9orf72 dipeptide repeat proteins identifies molecular chaperones as modifiers of poly-GA aggregation. Acta Neuropathol Commun 10: 22

Lorenzo G, 2013. Cell senescence : Methods and Protocols, Methods in Molecular Biology.

Maor-Nof M, Shipony Z, Lopez-Gonzalez R, Nakayama L, Zhang YJ, Couthouis J, Blum JA, Castruita PA, Linares GR, Ruan K et al, 2021. p53 is a central regulator driving neurodegeneration caused by C9orf72 poly(PR), Cell. Elsevier, pp. 689–708.e620.

Marine JC, Lozano G (2010) Mdm2-mediated ubiquitylation: p53 and beyond. Cell Death Differ 17: 93–102

Martínez-Cué C, Rueda N (2020) Cellular Senescence in Neurodegenerative Diseases. Frontiers in Cellular Neuroscience 14

Matias I, Diniz LP, Damico IV, Araujo APB, Neves LDS, Vargas G, Leite REP, Suemoto CK, Nitrini R, Jacob-Filho W et al (2022) Loss of lamin-B1 and defective nuclear morphology are hallmarks of astrocyte senescence in vitro and in the aging human hippocampus. Aging Cell 21: e13521

Mitra J, Guerrero EN, Hegde PM, Liachko NF, Wang H, Vasquez V (2019) Motor neuron disease-associated loss of nuclear TDP-43 is linked to DNA double-strand break repair defects. 1–10

Mizielinska S, Ridler CE, Balendra R, Thoeng A, Woodling NS, Grässer FA, Plagnol V, Lashley T, Partridge L, Isaacs AM, 2017. Bidirectional nucleolar dysfunction in C9orf72 frontotemporal lobar degeneration, Acta neuropathologica communications. Acta Neuropathologica Communications, p. 29.

Morrison RS, Kinoshita Y, 2000. The role of p53 in neuronal cell death, Cell Death and Differentiation. pp. 868–879.

Ohashi M, Korsakova E, Allen D, Lee P, Fu K, Vargas BS, Cinkornpumin J, Salas C, Park JC, Germanguz I et al (2018) Loss of MECP2 Leads to Activation of P53 and Neuronal Senescence. Stem Cell Reports 10: 1453–1463

Parlato R, Liss B, 2014. How Parkinson’s disease meets nucleolar stress, Biochimica et Biophysica Acta - Molecular Basis of Disease. Elsevier B.V., pp. 791–797.

Piol D, Tosatto L, Zuccaro E, Anderson EN, Falconieri A, Polanco MJ, Marchioretti C, Lia F, White J, Bregolin E et al (2023) Antagonistic effect of cyclin-dependent kinases and a calcium-dependent phosphatase on polyglutamine-expanded androgen receptor toxic gain of function. Sci Adv 9: eade1694

Ramesh N, Daley EL, Gleixner AM, Mann JR, Kour S, Mawrie D, Anderson EN, Kofler J, Donnelly CJ, Kiskinis E et al (2020a) RNA dependent suppression of C9orf72 ALS/FTD associated neurodegeneration by Matrin-3. Acta Neuropathol Commun 8: 177

Ramesh N, Kour S, Anderson EN, Rajasundaram D, Pandey UB (2020b) RNA-recognition motif in Matrin-3 mediates neurodegeneration through interaction with hnRNPM. Acta Neuropathol Commun 8: 138

Renton AE, Majounie E, Waite A, Simón-Sánchez J, Rollinson S, Gibbs JR, Schymick JC, Laaksovirta H, van Swieten JC, Myllykangas L et al (2011) A hexanucleotide repeat expansion in C9ORF72 is the cause of chromosome 9p21-linked ALS-FTD. Neuron 72: 257–268

Saez-Atienzar S, Masliah E (2020) Cellular senescence and Alzheimer disease: the egg and the chicken scenario. Nature Reviews Neuroscience

Salminen A, Kauppinen A, Kaarniranta K (2012) Emerging role of NF-kappaB signaling in the induction of senescence-associated secretory phenotype (SASP). Cell Signal 24: 835–845

Sasaki M, Kawahara K, Nishio M, Mimori K, Kogo R, Hamada K, Itoh B, Wang J, Komatsu Y, Yang YR et al (2011) Regulation of the MDM2-P53 pathway and tumor growth by PICT1 via nucleolar RPL11. Nat Med 17: 944–951

Shaker MR, Aguado J, Chaggar HK, Wolvetang EJ (2021) Klotho inhibits neuronal senescence in human brain organoids. NPJ Aging Mech Dis 7: 18

Trias E, Beilby PR, Kovacs M, Ibarburu S, Varela V, Barreto-Núñez R, Bradford SC, Beckman JS, Barbeito L, 2019. Emergence of microglia bearing senescence markers during paralysis progression in a rat model of inherited ALS, Frontiers in Aging Neuroscience. pp. 1–14.

Van Deursen JM, 2014. The role of senescent cells in ageing, Nature. Nature Publishing Group, pp. 439–446.

Vazquez-Villasenor I, Garwood CJ, Heath PR, Simpson JE, Ince PG, Wharton SB (2020) Expression of p16 and p21 in the frontal association cortex of ALS/MND brains suggests neuronal cell cycle dysregulation and astrocyte senescence in early stages of the disease. Neuropathol Appl Neurobiol 46: 171–185

Wang H, Hegde ML (2019) New Mechanisms of DNA Repair Defects in Fused in Sarcoma-Associated Neurodegeneration: Stage Set for DNA Repair-Based Therapeutics? J Exp Neurosci 13: 1179069519856358

Wei Z, Chen XC, Song Y, Pan XD, Dai XM, Zhang J, Cui XL, Wu XL, Zhu YG (2016) Amyloid beta Protein Aggravates Neuronal Senescence and Cognitive Deficits in 5XFAD Mouse Model of Alzheimer’s Disease. Chin Med J (Engl) 129: 1835–1844

Wen X, Tan W, Westergard T, Krishnamurthy K, Markandaiah SS, Shi Y, Lin S, Shneider NA, Monaghan J, Pandey UB et al (2014) Antisense proline-arginine RAN dipeptides linked to C9ORF72-ALS/FTD form toxic nuclear aggregates that initiate invitro and invivo neuronal death. Neuron 84: 1213–1225

Zhang Y-j, Guo L, Gonzales PK, Gendron TF, Wu Y, Jansen-west K, Raw ADO, Pickles SR, Prudencio M, Carlomagno Y et al (2019) Heterochromatin anomalies and double-stranded RNA accumulation underlie C9orf 72 poly(PR) toxicity. 2606

Zhang YJ, Gendron TF, Ebbert MTW, O’Raw AD, Yue M, Jansen-West K, Zhang X, Prudencio M, Chew J, Cook CN et al (2018) Poly(GR) impairs protein translation and stress granule dynamics in C9orf72-associated frontotemporal dementia and amyotrophic lateral sclerosis. Nature Medicine 24: 1136–1142

Zhang YJ, Jansen-West K, Xu YF, Gendron TF, Bieniek KF, Lin WL, Sasaguri H, Caulfield T, Hubbard J, Daughrity L et al (2014) Aggregation-prone c9FTD/ALS poly(GA) RAN-translated proteins cause neurotoxicity by inducing ER stress. Acta Neuropathologica 128: 505–524

